# End-to-end learning of multiple sequence alignments with differentiable Smith-Waterman

**DOI:** 10.1101/2021.10.23.465204

**Authors:** Samantha Petti, Nicholas Bhattacharya, Roshan Rao, Justas Dauparas, Neil Thomas, Juannan Zhou, Alexander M. Rush, Peter K. Koo, Sergey Ovchinnikov

## Abstract

Multiple Sequence Alignments (MSAs) of homologous sequences contain information on structural and functional constraints and their evolutionary histories. Despite their importance for many downstream tasks, such as structure prediction, MSA generation is often treated as a separate pre-processing step, without any guidance from the application it will be used for. Here, we implement a smooth and differentiable version of the Smith-Waterman pairwise alignment algorithm that enables jointly learning an MSA and a downstream machine learning system in an end-to-end fashion. To demonstrate its utility, we introduce SMURF (Smooth Markov Unaligned Random Field), a new method that jointly learns an alignment and the parameters of a Markov Random Field for unsupervised contact prediction. We find that SMURF learns MSAs that mildly improve contact prediction on a diverse set of protein and RNA families. As a proof of concept, we demonstrate that by connecting our differentiable alignment module to AlphaFold and maximizing predicted confidence, we can learn MSAs that improve structure predictions over the initial MSAs. Interestingly, the alignments that improve AlphaFold predictions are self-inconsistent and can be viewed as adversarial. This work highlights the potential of differentiable dynamic programming to improve neural network pipelines that rely on an alignment and the potential dangers of relying on black-box methods for optimizing predictions of protein sequences.

---

Multiple Sequence Alignments (MSAs) are commonly used in biology to model evolutionary relationships and the structural/functional constraints within families of proteins and RNA. MSAs are a critical component of the latest contact [6, 28, 41] and protein structure prediction pipelines [5, 30]. Moreover, they are used for predicting the functional effects of mutations [19, 20, 27, 59], phylogenetic inference [18] and rational protein design [21, 37, 53, 61]. Creating alignments, however, is a challenging problem. Standard approaches use heuristics for penalizing substitutions and gaps and do not take into account the effects of contextual interactions [57] or long-range dependencies. For example, these local approaches struggle when aligning large numbers of diverse sequences, and additional measures (such as the introduction of external guide Hidden Markov Models, HMMs) must be introduced to obtain reasonable alignments [55]. Finally, each alignment method has a number of hyperparameters which are often chosen on an application-specific basis. This suggests that computational methods that input an MSA could be improved by jointly learning the MSA and training the method.

Here we introduce *Learned Alignment Module* (LAM), which is a fully differentiable module for constructing MSAs and hence can be trained in conjunction with another differentiable downstream task. Building upon the generalized framework for differentiable dynamic programming developed in [38], LAM employs a smooth and differentiable version of the Smith-Waterman algorithm. Whereas the classic implementation of the Smith-Waterman algorithm outputs a pairwise alignment between two sequences that maximizes an alignment score [56], the smooth version outputs a distribution over alignments. This smoothness is crucial to: (i) make the algorithm differentiable and therefore applicable in end-to-end neural network pipelines, and (ii) allow the method to consider multiple hypothesized alignments simultaneously, which we believe to be a beneficial feature early in training.

We demonstrate the utility of LAM with two differentiable pipelines. First, we design an unsupervised contact prediction method that jointly learns an alignment and the parameters of a Markov Random Field (MRF) for RNA and protein, which we use to infer better structure-based contact maps. Next, we connect our differentiable alignment method to AlphaFold to jointly infer an alignment that improves its prediction of protein structures [30]. Our main contributions are as follows:

1. We implemented a smooth and differentiable version of the Smith-Waterman algorithm for local pairwise alignment in JAX [10]. Our implementation includes options for an affine gap penalty, a temperature parameter that controls the relaxation from the highest scoring path (i.e. smoothness), and both global and local alignment settings. Our code is freely available and can be applied in any end-to-end neural network pipeline written in JAX, TensorFlow [1] or via DLPack in PyTorch [50]. Moreover, we give a self-contained description of our implementation and its mathematical underpinnings, providing a template for future implementations in other languages.
2. We introduced the Learned Alignment Module (LAM), a fully differentiable module for constructing MSAs that is trained in conjunction with a downstream task. For each input sequence, a convolutional architecture produces a matrix of match scores between the sequence and a reference sequence. Unlike a substitution matrix typically input to Smith-Waterman, these scores account for the local *k*-mer context of each residue. Next we apply our smooth Smith-Waterman implementation to these similarity matrices to align each sequence to the reference, yielding an MSA (Fig. 1).
3. We designed a method called *Smooth Markov Unaligned Random Field* (SMURF) that takes as input unaligned sequences and jointly learns an MSA (via LAM) and MRF parameters. These parameters can then be used for contact prediction. We show that SMURF outperforms GREMLIN, when trained with the same objective, for protein and RNA contact prediction on a diverse set of families.
4. To demonstrate the utility of a differentiable alignment layer, we modify AlphaFold [30], replacing the input MSA with the output of LAM. For a given set of unaligned, related protein sequences, we backprop through AlphaFold to update the parameters of LAM, maximizing AlphaFold’s predicted confidence. Doing so results in learned MSAs that improve the structure prediction over our initial input MSA for 3 out of 4 targets. Despite the improved structure predictions, we find that the MSAs learned by the LAM may be adversarial as indicated by their self-inconsistency. This finding raises questions about how AlphaFold uses the input MSA to make its predictions.

**FIG. 1:**
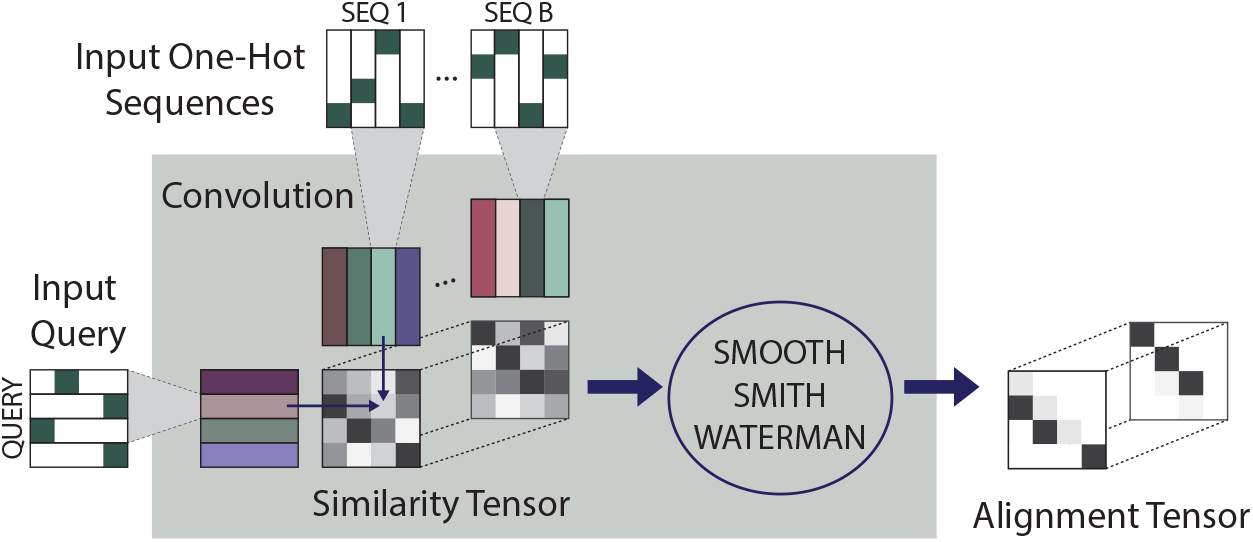
Learned alignment module (LAM). The residues of *B* sequences and a “query” sequence are mapped to vectors using a convolution. For each sequence *k*, an alignment score matrix *a* is computed by taking the dot products of the vectors representing the query sequence and the vectors representing sequence *k*. The similarity tensor is formed by concatenating these matrices, and then our differentiable implementation of smooth Smith-Waterman is applied to each similarity matrix in the tensor to produce an alignment. The resulting *B* smooth pairwise alignments (all aligned to the query sequence) are illustrated as the “Alignment Tensor.”

## Related work

### a. Differentiable Dynamic Programming in Natural Language Processing (NLP)

Differentiable dynamic programming algorithms are needed in order to model combinatorial structures in a way that allows backpropagation of gradients [8, 38, 62]. Such algorithms have been used in NLP to build neural models for parsing [16], grammar induction [33], speech [11], and more. Smooth relaxations of argmax and other non-differentiable functions can enable differentiation through dynamic programs. More generally, Mensch and Blondel leverage semirings to provide a unified framework for constructing differentiable operators from a general class of dynamic programming algorithms [38]. This work has been incorporated into the Torch-Struct library [52] to enable composition of automatic differentiation and neural network primitives, was recently implemented in Julia [58], and is the basis for our JAX implementation of smooth Smith-Waterman.

### b. Smooth and differentiable alignment in computational biology

Before end-to-end learning was common, computational biologists used pair HMMs to express probability distributions over pairwise alignments [15, 35, 40]. The forward algorithm applied to a pair HMM can be viewed as a smoothed version of Smith-Waterman. Later, a differentiable kernel-based method for alignment was introduced [54]. More recently, Morton et al. implemented a differentiable version of the Needleman-Wunsch algorithm for global pairwise alignment [43, 46]. Our implementation has several advantages: (i) vectorization makes our code faster (Fig. S1 and Supplementary Note S1 C), (ii) we implemented local alignment and an affine gap penalty (Supplementary Note S1 D), and (iii) due to the way gaps are parameterized, the output of [43] can not be interpreted as an expected alignment (Supplementary Note S1 B). Independent and concurrent work [36] uses a different formulation of differentiable Smith-Waterman involving Fenchel-Young loss.

### c. Language models, alignments, and MRFs

Previous work combining language model losses with alignment of biological sequences place the alignment layer at the end of the pipeline. Bepler et al. first pretrain a bidirectional RNN language model, then freeze this model and train a downstream model using a pseudo-alignment loss [7]. Similarly, Morton et al. use a pretrained language model to parametrize the the alignment scoring function [43]. Their loss, however, is purely supervised based on ground-truth structural alignments. Llinares-López et al. use differentiable Smith-Waterman with masked language modeling and supervised alignments to learn a scoring function dervived from transformer embeddings [36]. For RNA, a transformer embedding has been trained jointly with a masked language modeling and structural alignment [2]. In contrast to all of these papers, our alignment layer is in the middle of the pipeline and is trained end-to-end with a task downstream of alignment.

Joint modeling of alignments and Potts models has been explored. Kinjo et al. [34] include insertions and deletions into a Potts model using techniques from statistical physics. Two other works infer HMM and/or Potts parameters through importance sampling [63] and message passing [44], with the goal of designing generative classifiers for protein homology search.

## RESULTS

### Smooth Smith-Waterman

Pairwise sequence alignment is the task of finding an alignment of two sequences with the highest score, where the score is the sum of the “match” scores for each pair of aligned residues and “gap” penalties for residues that are unmatched. The Smith-Waterman algorithm is a dynamic programming algorithm that returns a path with the maximal score. A *smooth* version instead finds a probability distribution over paths in which higher scoring paths are more likely. Smoothness and differentiability can be achieved by replacing the max with logsumexp and argmax with softmax in the dynamic programming algorithm. We implemented a Smooth Smith-Waterman (SSW) formulation in which the probability that any pair of residues is aligned can be formulated as a derivative (see Methods). We use JAX due to its JIT (‘just in time’) compilation and automatic differentiation features [10].

Our speed benchmark indicates that our implementation is faster than the smooth Needleman-Wunsch implementation in [43] for both a forward pass as well as for the combined forward and backward passes, see Fig. S1. The latter is relevant when using the method in a neural network pipeline requiring backprogation. Moreover, comparison between a vectorized and naive version of our code shows that vectorization substantially reduces the runtime, see [64] and Supplementary Note S1 C.

Our SSW has four other features: temperature, affine gap, retrict turns, and global alignment. A *temperature* parameter governs the extent to which the distribution concentrated on the highest scoring alignments. In the *affine gap* mode, the first gap in a streak incurs an “open” gap penalty and all subsequent gaps incur an “extend” gap penalty. A *restrict turns* option corrects for the algorithm’s inherent bias towards alignments near the diagonal. We also implemented Needleman-Wunsch to output *global alignments* rather than local alignments. See Supplementary Note S1 D for additional details of SSW options.

### Learned Alignment Module (LAM)

The key to improving a Smith-Waterman alignment is finding the right input matrix of alignment scores *a* = (*a*_*ij*_)_*i≤𝓁x*,*j≤𝓁y*_. Typically, when Smith-Waterman is used for pairwise alignment the alignment score between positions *i* and *j, a*_*ij*_, is given by a BLOSUM or PAM score for the pair of residues *X*_*i*_ and *Y*_*j*_ [3, 13, 24]. This score reflects how likely it is for one amino acid to be substituted for another, but does not acknowledge the context of each residue in the sequence. For example, consider serine, an amino acid that is both small and hydrophilic. In a water-facing part of a protein, serine is more likely to be substituted for other hydrophilic amino acids. In other contexts, serine may only be substituted for other small amino acids due to the geometric constraints of the protein fold. Employing a scoring function with convolutions allows for local context to be considered.

Our proposed learned alignment module adaptively learns a context-dependent alignment score matrix *a*_*ij*_, performs an alignment based on this score matrix, all *in conjunction with a downstream machine learning task*. The value *a*_*ij*_ expresses the similarity between *X*_*i*_ in the context of *X*_*i−w*_, … *X*_*i*_, … *X*_*i*+*w*_ and *Y*_*j*_ in the context of *Y*_*j−w*_, … *Y*_*j*_, … *Y*_*j*+*w*_. We represent position *i* in sequence *X* as a vector 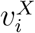 obtained by applying a convolutional layer of window size 2*w* + 1 to a one-hot encoding of *X*_*i*_ and its neighbors. The value *a*_*ij*_ in the similarity matrix that we input to Smith-Waterman is the dot product of the corresponding vectors, 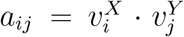. To construct an MSA from a reference and *B* other sequences, the LAM constructs a similarity matrix between each sequence and the reference, applies differentiable Smith-Waterman to each similarity matrix, and outputs an alignment of each sequence to the reference (which can be viewed as an MSA). See Fig. 1. Since process is entirely differentiable, we can plug the alignment produced by the LAM into a downstream module, compute a loss function, and train the whole pipeline end-to-end.

### Applying the LAM to contact prediction

GREMLIN is a probabilistic model of protein variation that uses the MSA of a protein family to estimate parameters of a MRF (see Methods), which in turn are used to predict contact maps [6, 17, 32, 49]. Since GREMLIN relies on an input MSA, one would expect that improved alignments would yield better contact prediction results. To test this, we designed a pipeline for training a GREMLIN-like model that inputs unaligned sequences and jointly learns the MSA and MRF parameters. We call our method **S**mooth **M**arkov **U**naligned **R**andom **F**ield or SMURF.

SMURF takes as input a family of unaligned sequences and learns both (i) the LAM convolutions and (ii) the parameters of the MRF that are, in turn, used to predict contacts. SMURF has two phases, each beginning with the LAM. First, BasicAlign learns LAM convolutions by minimizing the squared difference between each aligned sequence and the corresponding averaged MSA (Fig. S5). These convolutions are then used to initialize the LAM for the second training phase, TrainMRF, where a masked language modeling (MLM) objective is used to learn MRF parameters and update the convolutions (Fig. S6). We compare SMURF to GREMLIN trained with masked language modeling (MLM-GREMLIN) [9]. The architecture of MLM-GREMLIN is the similar to TrainMRF step of SMURF, except that a fixed alignment is input instead of a learned alignment computed by LAM.

We trained and evaluated our model on a diverse set of protein families, as described in Methods. To evaluate the accuracy of downstream contact prediction, we computed a standard metric used to summarize contact prediction accuracy, i.e. the area under the curve (AUC) for a plot of fraction of top *t* predicted contacts that are correct for *t* equals 1 up to *L*, where *L* is the length of the protein. Fig. 2a illustrates that SMURF mildly outperforms MLM-GREMLIN with a median AUC improvement of 0.007 across 193 protein families in the test set. To test whether SMURF requires a deep alignment with many sequences, we ran SMURF on protein families at most 128 sequences. The performance of SMURF and MLM-GREMLIN are comparable even for these families with relatively few sequences, with a median AUC improvement of 0.002 (Fig. S8).

**FIG. 2:**
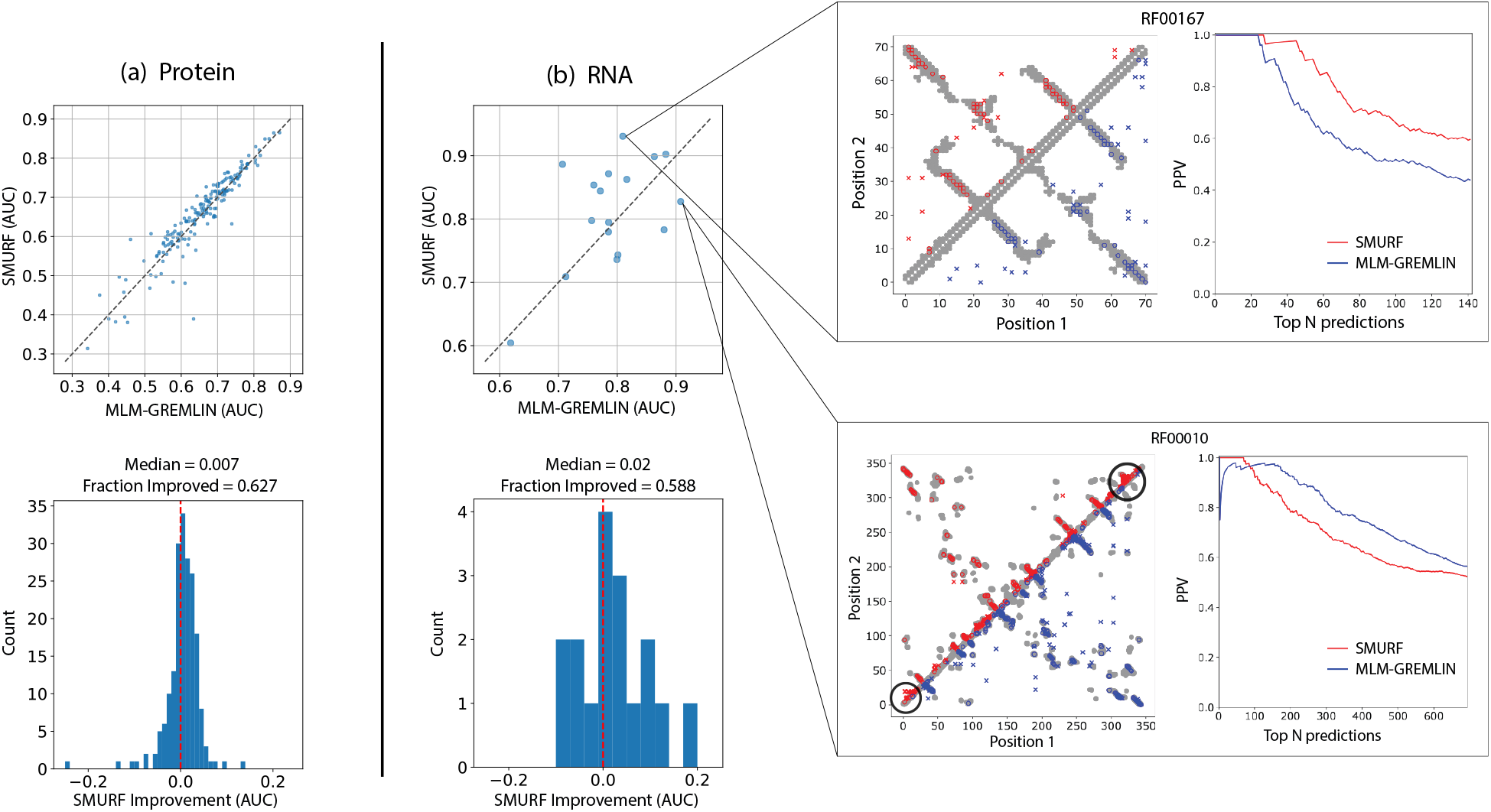
SMURF outperforms MLM-GREMLIN on (a) protein and (b) non-coding RNA. (Top) Scatter plots of the AUC of the top L predicted contacts for SMURF versus MLM-GREMLIN. (Bottom) Histograms of the difference in AUC between SMURF and MLM-GREMLIN. (Right) Comparison of contact predictions and the positive predictive value (PPV) for different numbers of top *N* predicted contacts, with *N* ranging from 0 to 2*L*, for SMURF (red) and MLM-GREMLIN (blue) for Rfam family RF00010 (Ribonuclease P.) and RF00167 (Purine riboswitch). Gray dots represent PDB-derived contacts, circles represent a true positive prediction, and x represents a false positive prediction. For contact predictions for RFAM00010, the black circles highlight a concentration of false positive predictions.

Next we sought to compare qualities of the MSAs learned through SMURF and MSAs fed into GREMLIN, which were generated with HHblits [57]. To quantify the consistency of the MSAs, we compared the BLOSUM scores [24] of all pairwise alignments extracted from our learned MSA to those extracted from the HHblits MSA. By this metric, we found that alignments learned by SMURF were more consistent than those from HHblits. Moreover, we observed a slightly positive correlation between increased consistency and contact prediction improvement (Fig. S7, left). We also found that SMURF alignments tend to have more positions aligned to the query (Fig. S7, right). We hypothesize that this is because our MRF does not have a mechanism to intelligently guess the identity of residues that are insertions with respect to the query sequence (the guess is uniform, see Methods).

Next, we applied SMURF to 17 non-coding RNA families from Rfam [31] that had a corresponding structure in PDB (see Methods). Due to the relatively small number of RNAs with known 3D structures, we employed SMURF using the hyperparameters optimized for proteins; fine-tuning SMURF for RNA could improve performance. Overall, we observe that SMURF outperforms MLM-GREMLIN with a median AUC improvement of 0.02 (Fig. 2b). In Supplementary Note S2, we further discuss the RNA contact predictions illustrated in Fig. 2b and the SMURF predictions for the three most and least improved protein families (Figs. S9 and S10). We hypothesize that SMURF generates fewer false positive predictions in seemingly random locations because the LAM finds better alignments.

Finally, we performed an ablation study on SMURF (Fig. S11). We found that replacing smooth Smith-Waterman with a differentiable “pseudo-alignment” procedure, similar to [7], degraded performance substantially. Skipping BasicAlign also degraded performance, thus indicating the importance of the initial convolutions found in BasicAlign.

### Using backprop through AlphaFold to learn alignments with LAM

As a proof of concept, we selected four CASP14 domains where the structure prediction quality from AlphaFold was especially sensitive to how the MSA was constructed. We reasoned that the quality was poor due to issues in the MSA and by realigning the sequences using AlphaFold’s confidence metrics we may be able to improve on the prediction quality. For each of the four selected CASP targets, separate LAM parameters were fit to maximize AlphaFold’s predicted confidence metrics (see Methods). We repeated this 180 times for each target (varying the learning rates, random seeds, and smoothness of the alignment), and then selected the learned MSA corresponding to the most confident AlphaFold (AF) prediction as measured by AF’s predicted local Distance Difference Test (pLDDT). For all targets, AF reported higher confidence in the prediction from our learned MSA as compared to the prediction from an MSA with the same sequences generated by MMSeqs2 as implemented in ColabFold [39]. However only 3 of the 4 targets showed an improvement in the structure prediction, as measured by the RMSD (root-mean-squared-distance) to native structure (see Figs. 3 and 4).

**FIG. 3:**
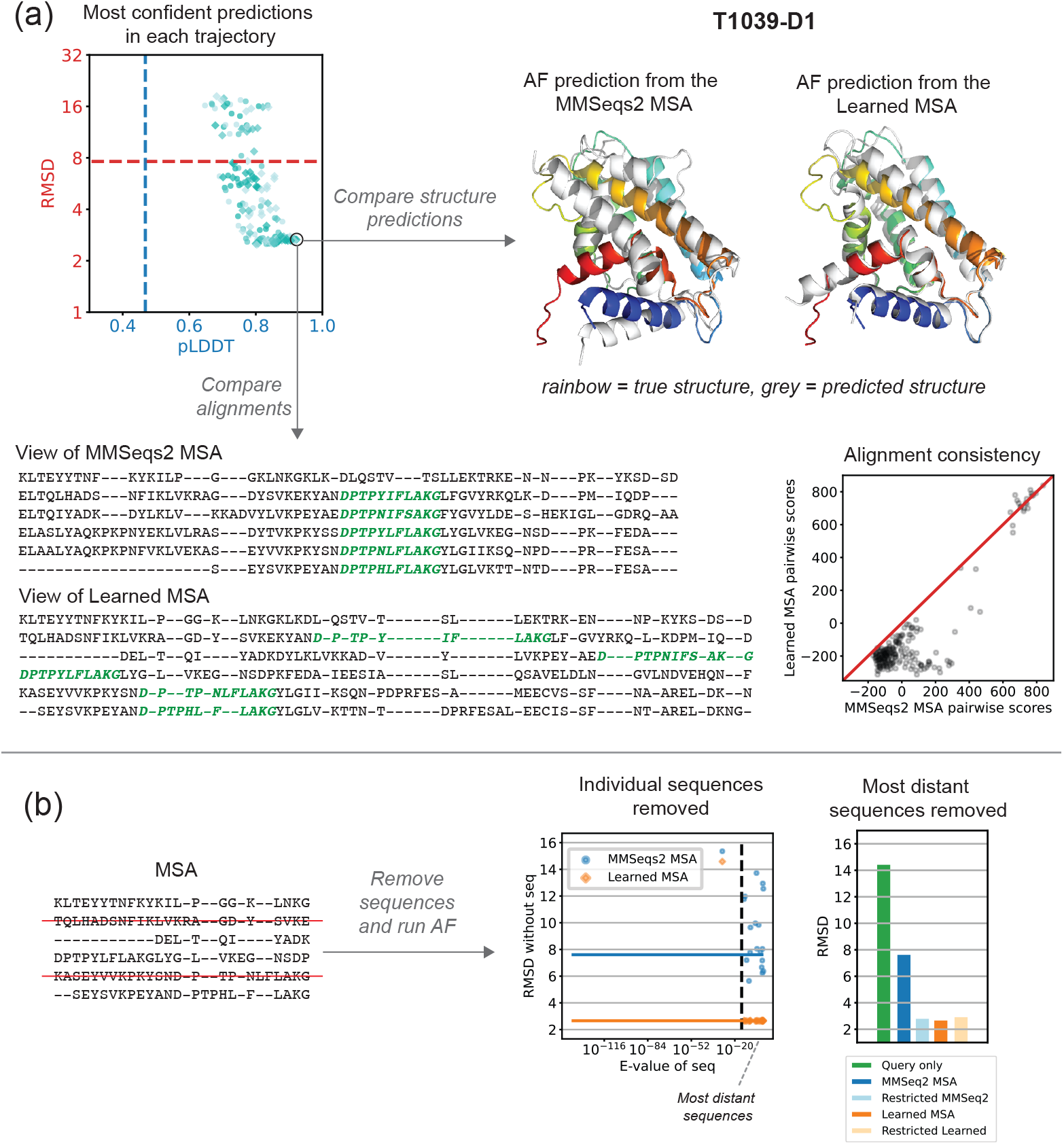
Learned MSA results in improved structure prediction, but a worse alignment for T1039-D1. (a) The scatter plot shows the pLDDT and RMSD for the most confident point in each trajectory. The marker color indicates the learning rate (10^*−*2^, 10^*−*3^, 10^*−*4^, lighest to darkest) and the shape indicates whether cooling was used (circle = no cooling, square = cooling). The dotted lines show the pLDDT and RMSD of the prediction using the MSA from MMseqs2. We selected the circled point maximizing the confidence (pLDDT) as our “Learned MSA.” The native structure is rainbow colored, and the predictions are overlaid in grey. The view of our Learned MSA illustrates the inconsistent alignment of a conserved motif (green) that is aligned accurately in the MMSeqs2 MSA. The scatter plot shows that the pairwise alignment scores for pairs extracted from the Learned MSA are much lower than the scores for pairs extracted from the MMSeqs2 MSA. (b) Change in RMSD when individual sequences are removed from the MSA (left) or a group of distant sequences is removed (right).

**FIG. 4:**
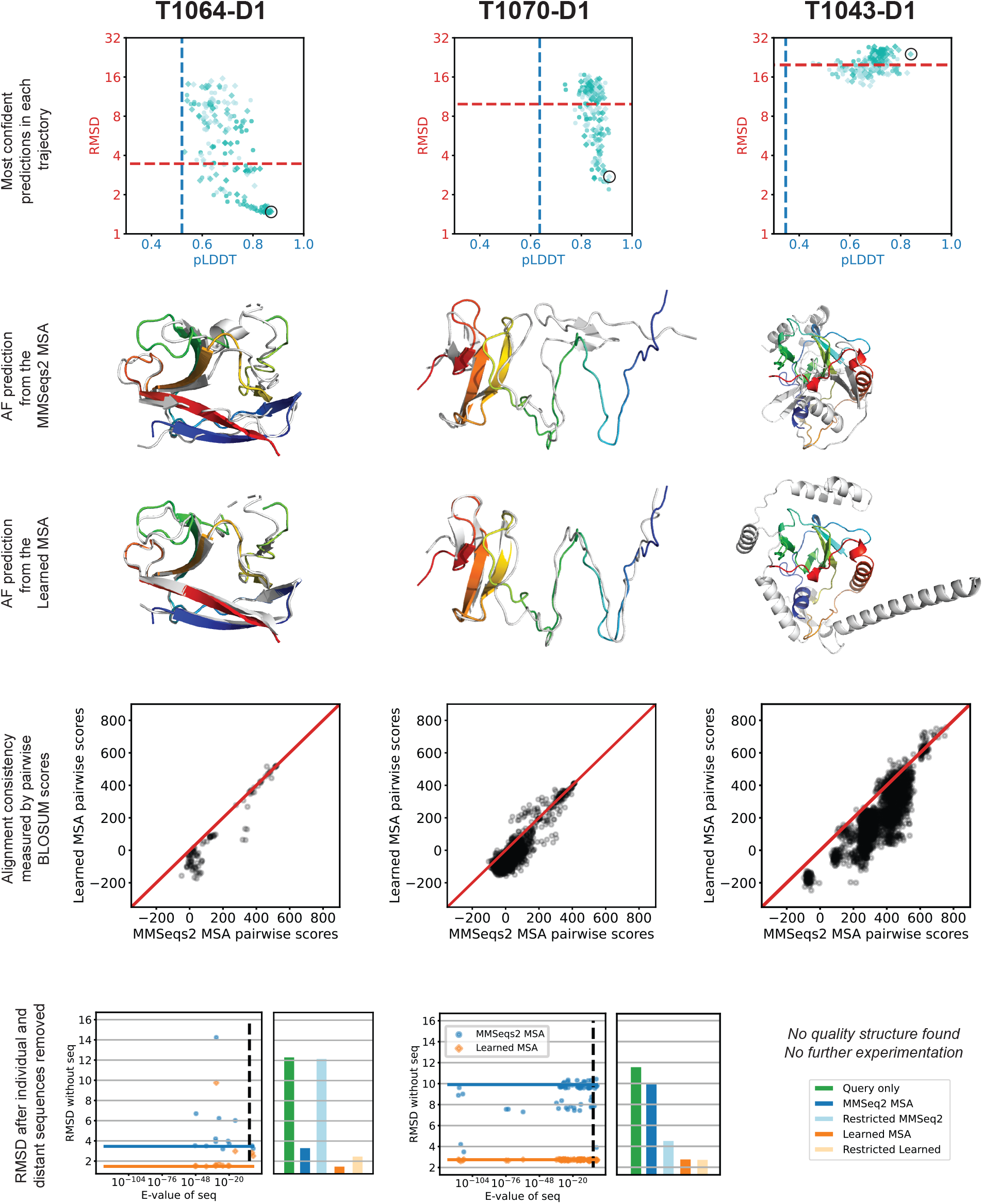
Learned MSA and structure predictions for three additional targets. The plots are analogous to those in Fig. 3. An improved structure was found for T1064-D1 and T1070-D1, but not T1043-D1. The MSAs learned for each target were less consistent than their MMSeqs2 counterparts.

Next we compared the learned MSAs that led to better structure predictions to the MMSeqs2 MSAs. Strikingly, we found our learned MSAs to be very low-quality. Fig. 3a illustrates a conserved motif that is consistently aligned in the MMSeqs2 MSA yet completely scattered in our learned MSA. To quantify the consistency of the MSAs, we compared the BLOSUM scores [24] of all pairwise alignments extracted from our learned MSAs to those extracted from the MMSeqs2 MSA. Indeed, the learned MSAs contain much lower scoring pairwise alignments than those of MMSeqs2 MSAs, indicating far less consistency (Figs. 3a and 4), which is the opposite trend we observed for MSAs learned by SMURF. Thus, unlike optimizing the MRF in SMURF, optimizing the confidence of AF predictions does not yield consistent alignments with LAM.

We explored a simple explanation for how low-quality alignments could yield improved structure predictions; perhaps AF uses its axial-like attention to consider only a subset of sequences, and the poor alignments by the other sequences isn’t important or could further disqualify those sequences from being attended to. To investigate this, we evaluated how sensitive the AF predictions are to the inclusion of each individual sequence (Figs. 3b and 4). Surprisingly, the prediction accuracy can be incredibly sensitive to the removal of a single sequence, especially for MMSeqs2 MSAs.

Next, we considered the effect of removing subsets of more distant sequences. The MMSeqs2 MSAs were constructed with a lenient E-value threshold of 10, which may introduce sequences in the MSA that are not true homologs. For targets T1064-D1 and T1070-D1, we removed all sequences with an E-value greater than 10^*−*3^. The target T1064-D1 has two sequences above this threshold (E-values 1.4 and 0.16) that almost certainly are not homologs of the query. (E-value, defined as P-value multiplied by the size of database, indicates the how many matches with detected similarity are expected to occur by chance alone.) While removing either individually does not substantially change the accuracy of the prediction, removing both worsens the prediction with the MMSeqs2 MSA significantly (RMSD 3.46 to 12.11) and worsens the prediction with our learned MSA mildly (RMSD 1.47 to 2.48). In T1070-D1 we realized the opposite outcome; removing the sequences with E-value at least 10^*−*3^ greatly improved the prediction with the MMSeqs2 MSA (RMSD 9.91 to 4.51) and slightly improved the prediction with our learned MSA (RMSD 2.75 to 2.70). Noting the influence of the closest homolog (E-value 6.1 *×* 10^*−*30^) on predictions for T1039-D1, we defined most distant sequences for this target as those with E-value greater than 10^*−*15^, leaving only the closest homolog. Restricting to the query and this single homolog improved the MMSeqs2 prediction substantially (RMSD 7.62 to 2.79), bringing it on on par with the prediction from our learned MSA on the full set of sequences (RMSD 2.66). The inclusion of this single close homolog is vital; the RMSD of the prediction for the query sequence alone is 11.56.

Finally, we repeated our optimization experiment after removing the distant sequences (Fig. S13a). We found that the most confident MSAs learned without the distant sequences tended to yield predictions with similar RMSD to the predictions from the most confident MSAs learned on the full set of sequences. (See orange and purple bars in Fig. S13b). We also investigated whether it was easier or harder to obtain “near optimal” structure prediction (having an RMSD of 1.25 times the RMSD of the prediction of the learned MSA on the full set) with the restricted set of sequences as compared to the full set. For T1064-D1 our optimization scheme found “near optimal” structures more often with the set of sequences that includes the distant sequences. The opposite was the case for T1039-D1, and there was no strong difference for T1070-D1 (Fig. S13b).

## DISCUSSION

In this work we explored the composition of alignment in a pipeline that can be trained end-to-end without usage of any existing alignment software or ground-truth alignments. With SMURF, we trained alignments jointly with a well-understood MRF contact prediction approach and found mild improvement in accuracy using learned MSAs that were consistent and reasonable. When we instead optimized with AlphaFold’s confidence metrics, we found low-quality MSAs that yielded improved structure predictions. This suggests that in order to learn high-quality alignments in the context of another machine learning task, the task must require high-quality alignments, which we discovered is not the case for structure prediction with AlphaFold. Perhaps by changing our objective function to also penalize self-inconsistent alignments, we could learn more reasonable MSAs while still improving AlphaFold predictions. Our work both establishes the feasibility of pipelines which jointly learn alignments in conjunction with downstream machine learning systems and highlights the possibility of unexpectedly learning odd alignments when it is not well-understood how exactly the downstream task uses alignments.

While our findings that low-quality, self-inconsistent MSAs can yield improved AlphaFold predictions and that AlphaFold predictions may be quite sensitive to the inclusion of particular sequences may seem paradoxical, these observations reflect behaviors found across deep learning systems. It is well-known that deep neural networks are not robust to adversarial noise [60]. Experiments that use an image recognition neural network to optimize an input image so that the image is confidently classified into a particular category will not necessarily yield human recognizable image of the category [42, 47]. Studying adversarial examples has been one approach to trying to understand how neural networks form predictions [23, 25, 42]. Our differentiable alignment module could be used with AlphaFold to identify a range of alignments that yield a particular prediction. Studying these alignments could provide insight on which aspects of an alignment are used by AlphaFold to make its prediction.

Our smooth Smith-Waterman implementation is designed to be usable and efficient, and we hope it will enable experimentation with alignment modules in other applications of machine learning to biological sequences. There is ample opportunity for future work to systematically compare architectures for the scoring function in smooth Smith-Waterman. The use of convolutions led to relatively simple training dynamics, but other inductive biases induced by recurrent networks, attention mechanisms, or hand-crafted architectures could capture other signal important for alignment scoring. We also hope that the use of these more powerful scoring functions enables applications in remote homology search, structure prediction, or studies of protein evolution.

Besides MSAs, there are numerous other discrete structures essential to analysis of biological sequences. These include Probabilistic Context Free Grammars used to model RNA Secondary Structure [45] and Phylogenetic Trees used to model evolution. Designing differentiable layers that model meaningful combinatorial latent structure in evolution and biophysics is an exciting avenue for further work in machine learning and biology.

## METHODS

Our code and a detailed description of the data we used is available at: https://github.com/spetti/SMURF.

### Smooth and differentiable Smith-Waterman

Pairwise sequence alignment can be formulated as the task of finding the highest scoring path through a directed graph in which edges correspond to an alignment of two particular residues or to a gap. The edge weights are match scores for the corresponding residues or the gap penalty, and the score of the path is the sum of the edge weights. The Smith-Waterman algorithm is a dynamic programming algorithm that returns a path with the maximal score. A *smooth* version instead finds a probability distribution over paths in which higher scoring paths are more likely. We describe a Smooth Smith-Waterman formulation in which the probability that any pair of residues is aligned can be formulated as a derivative.

Fig. 5 illustrates an alignment graph. For sequences *x*_1_, *x*_2_, … *x*_*𝓁x*_ and *y*_1_, *y*_2_, …, *y*_*𝓁y*_, the vertex set contains grid vertices *v*_*ij*_ for 0 *≤ i ≤ 𝓁*_*x*_ and 0 *≤ j ≤ 𝓁*_*y*_, a source *s*, and a sink *t*. The directed edges are defined so that each path from *s* to *t* corresponds to a local alignment of the sequences. The table below describes the definitions, meanings, and weights of the edges.

**FIG. 5:**
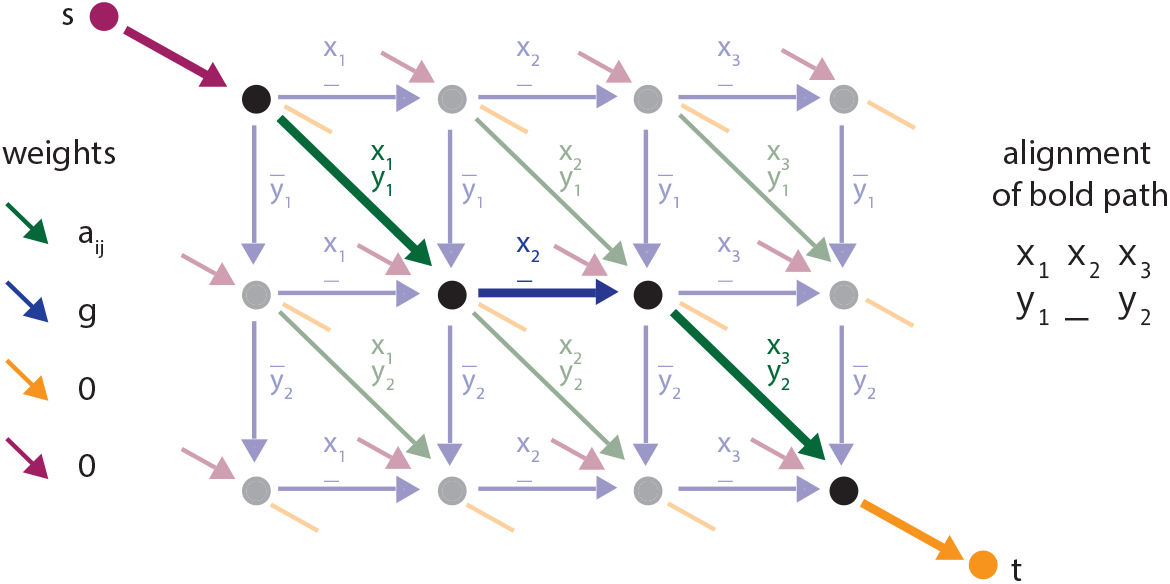
The alignment graph for sequences *X* = *x*_1_*x*_2_*x*_3_ **and** *Y* = *y*_1_*y*_2_. Edge labels describe the corresponding aligned pair, and colors indicate the weights. All red edges start at the source *s*, and all orange edges end at the sink *t*. The bold path corresponds to the alignment of *X* and *Y* written on the right.

| Edge | Meaning | Weight |
| --- | --- | --- |
| $v_{i-1,j-1} \rightarrow v_{i,j}$ | $x_i$ and $y_j$ are aligned | $x_i, y_j$ alignment score $a_{ij}$ |
| $v_{i,j-1} \rightarrow v_{i,j}$ | $y_j$ is aligned with the gap character - | gap penalty $g$ |
| $v_{i-1,j} \rightarrow v_{i,j}$ | $x_i$ is aligned with the gap character - | gap penalty $g$ |
| $s \rightarrow v_{i,j}$ | $x_k$ for $k \leq i$ and $y_k$ for $k \leq j$ are excluded | 0 |
| $v_{i,j} \rightarrow t$ | $x_k$ for $k > i$ and $y_k$ for $k > j$ are excluded | 0 |

The Smith-Waterman algorithm iteratively computes the highest score of a path ending at each vertex and returns the highest scoring path ending at *t*. Let *w*(*u → v*) denote the weight of the edge *u → v*, and let *N* ^*−*^(*v*) = *{u* | *u → v* is an edge*}* denote the incoming neighbors of *v*. Let *f* (*v*) be the value of the highest scoring path from *s* to *v*. Taking *f* (*s*) = 0, we compute

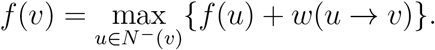

For grid vertices this simplifies to

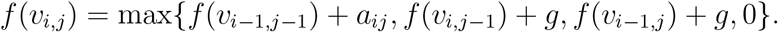

A path with the highest score is computed by starting at the sink *t* and tracing backward along the edges that achieve the maxima. (For further explanation see Chapter 2 of [15] or [56]).

Following the general differentiable dynamic programming framework introduced in [38], we implement a smoothed version of Smith-Waterman. We compute a smoothed version of the function *f*, which we denote *f* ^*S*^, by replacing the max with logsumexp. We again take *f* ^*S*^(*s*) = 0, and define

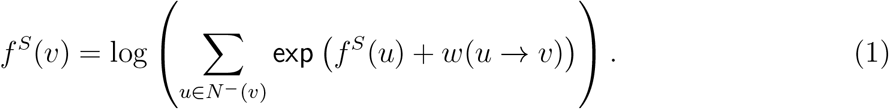

We use these smoothed scores and the edge weights to define a probability distribution over paths in *G*, or equivalently local alignments.

#### Definition 1.

*Given an alignment graph G* = (*E, V*), *define a random walk starting at vertex t that traverses edges of G in reverse direction according to transitions probabilities*

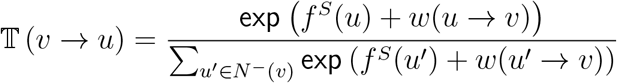

*and ends at the absorbing vertex s. Let µ*_*G*_ *be the probability distribution over local alignments in which the probability of an alignment A is equal to the probability that the random walk follows the reverse of the path in G corresponding to A*.

Under the distribution *µ*_*G*_, the probability that residues *x*_*i*_ and *y*_*j*_ are aligned can be formulated as a derivative. Mensch and Blondel describe this relationship in generality for differentiable dynamic programming on directed acyclic graphs [38]. We state their result as it pertains to our context and provide a proof in our notation in Supplementary Section S1 A.

#### Proposition 1

(Proposition 3 of [38]). *Let G be an alignment graph and µ*_*G*_ *be the corresponding probability distribution over alignments. Then*

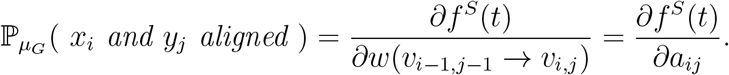

## GREMLIN

GREMLIN is a probabilistic model of protein variation that uses the MSA of a protein family to estimate parameters of a MRF of the form

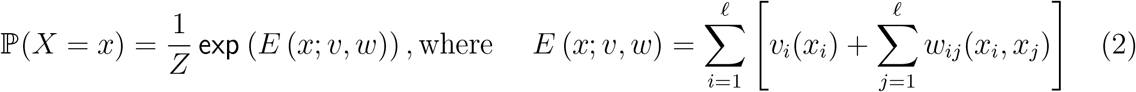

and *𝓁* is the number of columns in the MSA, *v*_*i*_ represents the amino acid propensities for position *i, w*_*ij*_ is the pairwise interaction matrix for positions *i* and *j*, and *Z* is the partition function (the value *E*(*·* ; *v, w*) summed over all sequences *x*). Typically the model is trained by maximizing the pseudo-likelihood of observing all sequences in the alignment [6, 17, 32, 49]. Here we follow the approach of [9, 51] and use *Masked Language Modeling* (MLM) to find the parameters *w* and *v*. The pairwise terms *w*_*ij*_ can be used to predict contacts by reducing each matrix *w*_*ij*_ into a single value that indicates the extent to which positions *i* and *j* are coupled.

### Data selection for SMURF

For our analysis of SMURF on proteins, we used the MSAs and contact maps collected in [4]. For training and initial tests, we used a reduced redundancy subset of 383 families constructed in [12]. Each family has least 1K effective sequences, and there is no pair of families with an E-value greater that 1e-10, as computed by an HMM-HMM alignment [26]. A random 190 families were used as the training set to identify quality hyperparameters of the model. The remaining 193 families served as the test set and are represented in Figure 2a, with the exceptions of two outlier families 4X9JA (SMURF AUC = 0.0748, MLM-GREMLIN AUC = 0.0523) and 2YN5A (SMURF AUC = 0.135, MLM-GREMLIN AUC = 0.145). Figure S8 includes data from 99 families from [26] that have at most 128 sequences. A list of the families used in each setting is available in our GitHub repository.

For each non-coding RNA, we aligned the RNA sequence in the PDB along with the corresponding Rfam sequences to an appropriate Rfam covariance model using Infernal [45]. We then analyzed these sequences using the same procedure outlined for proteins. We evaluated the efficacy of the predicted contact maps using the PDB-derived contact map, where two nucleotides are classified as in contact if the minimum atomic distance is below 8 angstrom. A list of the families used is available in our GitHub repository.

### Details of SMURF

SMURF has two phases: BasicAlign and TrainMRF. Both begin with the learned alignment module (Figure 1), but they have different architectures and loss functions afterwards.

#### BasicAlign

Similarity matrices produced by randomly initialized convolutions will produce chaotic alignments that are difficult for the downstream MRF to learn from. The purpose of BasicAlign is to learn initial convolutions whose induced similarity matrices yield alignments with relatively homogeneous columns (see Figure S5). The input to BasicAlign is a random subset of sequences *𝒮* = {*S*^(1)^, … *S*^(*B*)^} in the protein family. A pairwise alignment between each sequence and the first sequence *S*^(1)^ is produced via the learned alignment module (as described in Figure 1). This set of alignments can be viewed as an MSA where each column of the MSA corresponds to a position in the first sequence. Averaging the MSA yields the distribution of residues in each column. Let *M*_*ix*_ be the fraction of sequences in *𝒮* with residue *x* aligned to position *i* of *S*^(1)^,

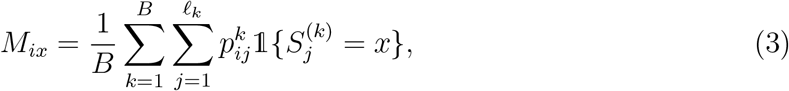

where *𝓁*_*k*_ is the length of *S*^(*k*)^ and 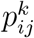 is the probability that position *i* of *S*^(1)^ is aligned to position *j* of *S*^(*k*)^ under the smooth Smith-Waterman alignment. (Note that Σ_*x*_ *M*_*ix*_ is less than one when there are sequences with a gap aligned to position *i* of *S*^(1)^.) The BasicAlign loss is computed by taking the squared difference between each aligned one-hot encoded sequence and the averaged MSA,

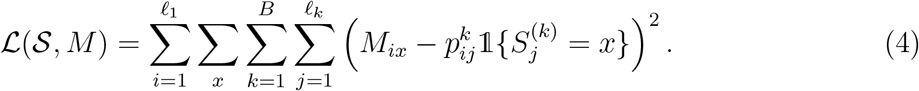

#### TrainMRF

In TrainMRF, masked language modeling is used to learn the MRF parameters and further adjust the alignment module convolutions (see Figure S6). The input to TrainMRF is a set of sequences drawn at random from the MSA, *𝒮* = {*S*^(1)^, … *S*^(*B*)^}. A random 15% of the residues of the input sequences are masked, and the masked sequences are aligned to the query via the learned alignment module (as described in Figure 1). The parameters for the alignment module are initialized from BasicAlign, and the query is initialized as the one-hot encoded reference sequence for the family.

The MRF has two sets of parameters: symmetric matrices *w*_*ij*_ *∈* ℝ^*A×A*^ for 1 *≤ i, j ≤ 𝓁*_*R*_ with *w*_*ij*_ = *w*_*ji*_ that correspond to pairwise interactions of the positions in the reference sequence and position-specific bias vectors *b*_*i*_ *∈* ℝ^*A*^ for 1 *≤ i ≤ 𝓁*_*R*_. Here *𝓁*_*R*_ denotes the length of the reference sequence, and *A* is the alphabet size (*A* = 20 for amino acids and *A* = 4 for nucleotides). Unlike traditional parameterizations of a MRF, we do not include gaps in our alphabet. Since our task is reconstructing masked positions in unaligned sequences, we have no need to predict gap characters.

After the sequences are aligned to the query, the infill distribution for each masked position is determined by the MRF parameters as follows. For a masked position *j* in sequence *k*, we define 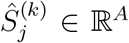 as the predicted probability distribution over residues at position *j* of sequence *S*^(*k*)^. Let 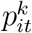 be the probability that position *t* of *S*^(*k*)^ is aligned to position *i* of the query under the smooth Smith-Waterman alignment, and let 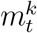 be the indicator that position *t* in sequence *S*^(*k*)^ was masked. To compute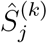, we first compute a score for each residue *x* that is equal to the expected value (under the smooth alignment) of the terms of the function *E*(*·* ; *b, w*) specific to position *j* or involving position *j* and an unmasked position. Then we compute the infill distribution by taking the softmax. Formally,

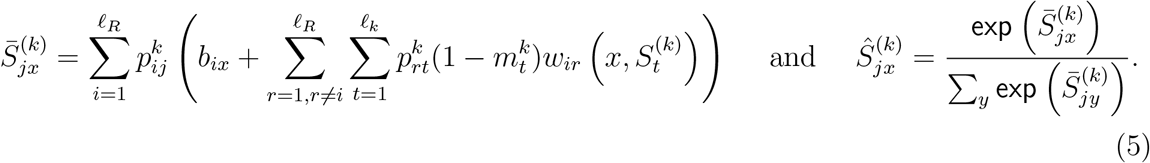

The infill distribution is an approximation of how likely each residue is to be present at position *j* in sequence *k* if position *j* were aligned to some position in the query sequence *S*^(1)^. The approximation considers the values of the linear terms *b* and the pairwise terms *w* corresponding only to unmasked positions. (In the case that position *j* in sequence *k* is almost certainly an insertion relative to the query sequence *S*^(1)^, i.e. 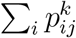 is small, our computation will likely provide a poor guess for the residue; in the extreme case where 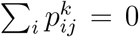 the infill distribution is uniform over the alphabet. Our model does not have a mechanism to learn the identities of residues that are insertions relative to the query sequence. Ultimately, this is not a concern since we do not use information about insertions to predict the contacts of the query sequence.)

We train the network using a cross entropy loss and *L*2 regularization on *w* and *b* with *λ* = .01

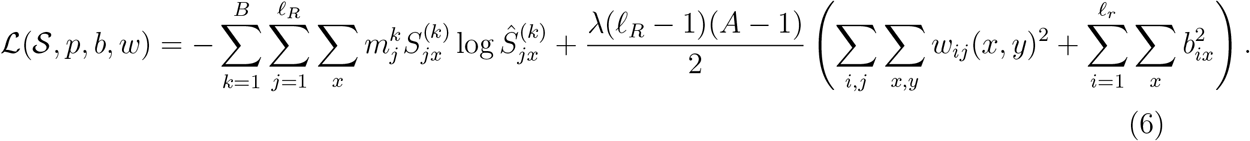

After each iteration, the query is updated to reflect the inferred MSA. Let *R* be the one-hot encoding of the reference sequence. We define *C*^*i*+1^ as a rolling weighted average of the MSAs learned through iteration *i* and *Q*^*i*^ as the query for iteration *i*,

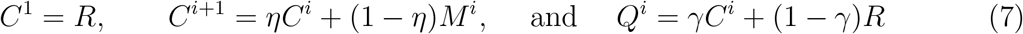

where *M*^*i*^ is the averaged MSA computed as described in Equation (3) from the sequences in iteration *i, η* = 0.90, and *γ* = 0.3. This process is illustrated by the light blue arrows in Figure S6. Preliminary results on the training set had suggested that updating the query in this manner improved results for some families. However, the ablation study on the test set does not suggest improvement (Fig. S11); further investigation is needed to determine the benefits changing the query between iterations.

Once training is complete, we use *w* to assign a contact prediction score between each pair of positions. The score *c*_*ij*_ measures the pairwise interaction between positions *i* and *j*, and 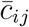 is score after applying APC correction [14],

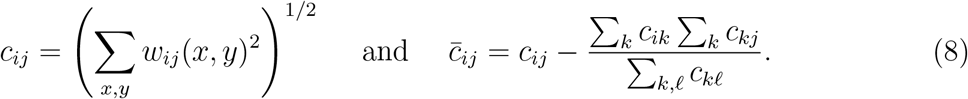

### SMURF hyperparameter selection

Throughout our hyperparameter search, we kept the following parameters constant: fraction of residues masked at 15%, number of convolution filters at 512, convolution window size at 18, regularization *λ* in Equation (6) at 0.01. Our hyperparameter search consisted of three stages. We initialized the gap penalty as *−*3 and allowed the network to learn a family-specific gap penalty.

1. First we ran a grid search with on all 190 families in the training set with learning rates *{*.05, 0.10, 0.15*}*, batch sizes *{*64, 128, 256*}*, and iterations *{*2000 BasicAlign */*1000 TrainMRF, 3000 BasicAlign */*3000 TrainMRF *}*. For comparison, we ran MLM-GREMLIN with the same range of learning rates and batch sizes and 3000 iterations. We found that batch size 64 and learning rate 0.05 performed best for MLM-GREMLIN.
2. Then we restricted to a smaller set of families to perform a more extensive hyperparameter search; we included the seven families where MLM-GREMLIN’s AUC was less than 0.45 (3AKBA, 3AWUA, 5BY4A, 4C6SA, 3OHEA, 3ERBA, 4F01A) and six families where SMURF consistently performed substantially worse than MLM-GREMLIN (1NNHA, 3AGYA, 4LXQA, 1COJA, 2D4XA, 4ONWA). We considered the following hyperparameter options: learning rates *{*.05, 0.10*}*, batch sizes *{*64, 128, 256*}*, iterations *{*2000 BasicAlign */*1000 TrainMRF, 2000 BasicAlign */*2000 TrainMRF, 3000 BasicAlign */*1000 TrainMRF *}*, MSA memory fraction *η ∈ {*0.90, 0.95*}*, and MSA query fraction *γ ∈ {*0.3, 0.5, 0.7*}*.
3. Based on the results of the above hyperparameter search on the select families, we performed a final hyperparameter search on the entire training set. We noticed that performance was better for larger batch sizes, but it was not always possible to run the large batch sizes on our 32 GB GPU for families with longer sequences. For our final hyperparameter search, we used the largest batch size of *{*64, 128, 256*}* that would fit in memory for each family. We set *η* = 0.90, *γ* = 0.3, and selected 3000 BasicAlign */*1000 TrainMRF iterations because these parameters lead to relatively strong results across the restricted set of families. Learning rate 0.10 outperformed learning rate 0.05 on the restricted set, but learning rate 0.05 generally outperformed learning rate 0.10 in the initial grid search on the full training set. We ran a final test with the aforementioned parameters and the two learning rates on the entire training set, and found that learning rate 0.05 was optimal overall.

We also ran 4000 iterations of MLM-GREMLIN with predetermined optimal parameters: learning rate 0.05 and batch size 64. We found very similar performance between 3000 and 4000 iterations of MLM-GREMLIN. We chose to compare SMURF to 4000 iterations of MLM-GREMLIN so that both methods were trained for 4000 iterations.

### Data selection for AlphaFold experiment

For our case study, the initial multiple sequence alignments (MSA) were obtained from MMseqs2 webserver as implemented in ColabFold [39]. After trimming the MSAs to their official domain definition, they were further filtered to reduce redundancy to 90 percent and to remove sequences that do not cover at least 75 percent of the domain length, using HHfilter [57]. Continuous domains under 200 in length, with at least 20 sequences, RMSD (root-mean-squared-distance) greater than 3 angstroms and the predicted LDDT (confidence metric) below 75, were selected for the experiment. We include one discontinuous targets T1064-D1 (SARS-CoV-2 ORF8 accessory protein) with only 16 sequences as an extra case study, as this was a particularly difficult CASP target that required manual MSA intervention, guided by pLDDT, to predict well [29]. The filtered MSAs were unaligned (gaps removed, deletions relative to query added back in) and padded to the max length.

### AlphaFold experiment details

We found that the AF predictions were particularly sensitive the the random mask used during evaluation (see Fig. S12). For this reason we omitted the mask during evaluation of the MMSeqs2 MSA and throughout our optimization procedure. For simplicity, we considered only one of the five AF models and did not give AF access to the length of insertions relative to the query during our optimization procedure. Our objective function sought to maximize the pLDDT and minimize the alignment error as returned by AF’s “model 3 ptm”. The AF predictions from the MMSeqs2 MSAs tended to have the overarching structure correct, but were incorrect on certain parts of the sequence. Our goal was for our optimization to correct the incorrect parts of the structure. For this reason we used the more stringent metric of RMSD (rather than the GDT measure of global structure) to evaluate the accuracy of our alignments.

When the number of sequences is low, we find the optimization to be especially sensitive to parameter initialization. To increase robustness, for each target 180 independent optimization trajectories with 100 iterations each were carried out using ADAM. Each trajectory is defined by a random seed, a learning rate (10^*−*2^, 10^*−*3^, 10^*−*4^) and whether a cooling scheme was used in Smith-Waterman (temperature 1.0 or temperature decreased linearly from 1.5 to 0.75 across the 100 iterations).

We thank Sean Eddy for pointing out the need for a restrict turns feature and for useful comments on a draft. We thank Jake VanderPlas for supplying JAX code that efficiently rotates a matrix (as in Figure S3b). We thank Tom Jones for helping us investigate the learned alignments from the AlphaFold experiment. Computational analyses were performed with assistance from the US National Institutes of Health Grant S10OD028632-01 and the Cannon cluster supported by the FAS Division of Science, Research Computing Group at Harvard University. SP was supported by the NSF-Simons Center for Mathematical and Statistical Analysis of Biology at Harvard (award #1764269). NB was supported in part by NIH grant R35-GM134922 and by the Exascale Computing Project (17-SC-20-SC), a collaborative effort of the U.S. Department of Energy Office of Science and the National Nuclear Security Administration. PKK was supported in part by the Simons Center for Quantitative Biology at Cold Spring Harbor Laboratory and the Developmental Funds from the Cancer Center Support Grant 5P30CA045508. SO is supported by NIH Grant DP5OD026389, NSF Grant MCB2032259 and the Moore–Simons Project on the Origin of the Eukaryotic Cell, Simons Foundation 735929LPI, https://doi.org/10.46714/735929LPI.

## Supplementary Information

### S1. SUPPLEMENTARY NOTE: SMOOTH SMITH-WATERMAN DETAILS AND FEATURES

#### A. Proof of the probabilistic interpretation of the gradient

**FIG. S1:**
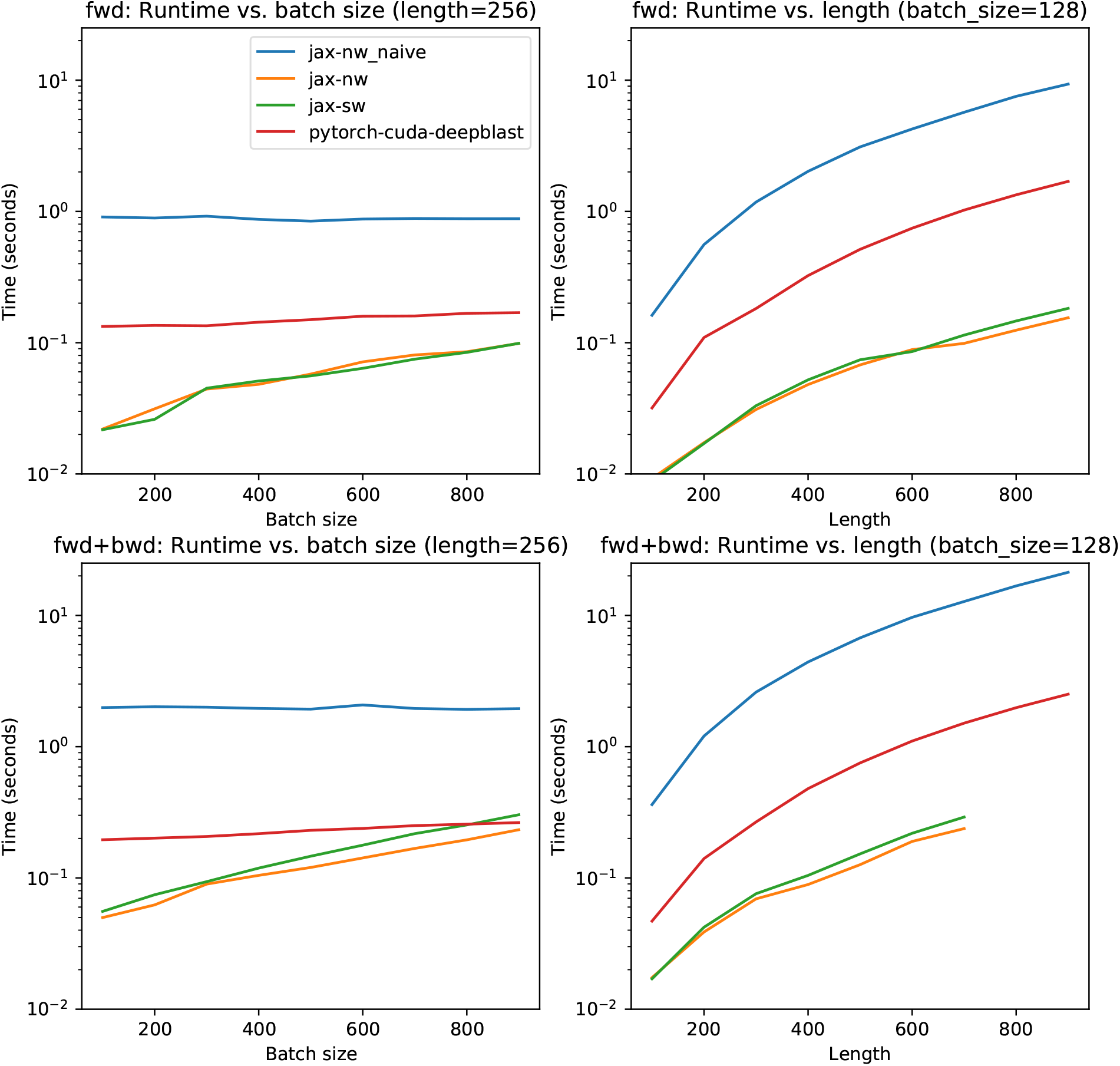
Runtime comparisons. We compare the runtimes of the Needleman-Wunsch implementation in [43] our JAX implementations of smooth Smith-Waterman (green), smooth Needleman-Wunsch (orange) and a naive non-vectorized Needleman-Wunsch (blue). Top plots report time for a forward pass, and the bottom plots report time for a forward and backward pass.

For completeness, we now repeat the proof of Proposition 1 given in [38] for the special case of Smooth Smith-Waterman. Proposition 1 gives a probabilistic interpretation of the gradient *f* ^*S*^(*t*) with respect to the edge weights *a*_*ij*_. We first give a probabilistic interpretation of the gradient *f* ^*S*^(*t*) with respect to the vertex scores *f* ^*S*^(*v*_*ij*_).

##### Proposition 2.

*Let G be an alignment graph. With respect to the random walk described in Definition 1*,

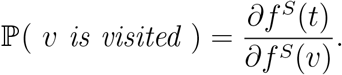

*Proof*. Let *N* ^+^(*v*) = *{u* | *v → u* is an edge in *G}* denote the outgoing neighborhood of *v*. Let *u*_1_, … *u*_*n*_ denote the vertices of *G* in a reverse topological order. We prove the statement by induction with respect to this order. Note *u*_1_ = *t*, and 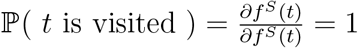. Assume that for all 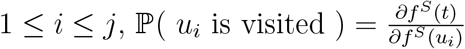. Observe

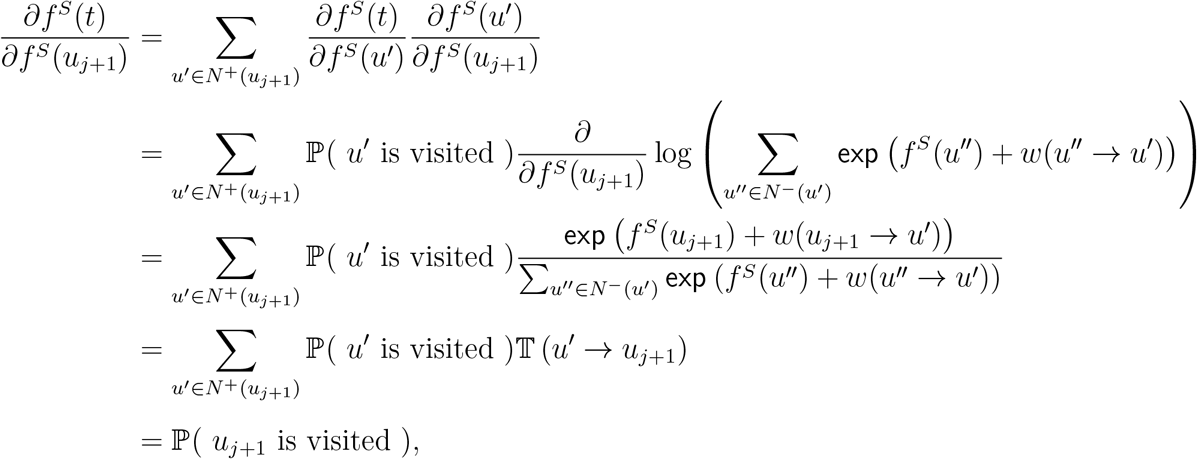

where in the second equality we apply the inductive hypothesis.

*Proof of Proposition 1*. It suffices to show that for each directed edge *u → v* in *G*

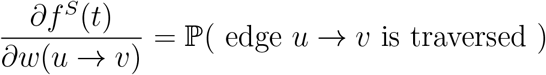

where the traversal occurs from *v* to *u* in the random walk. Observe

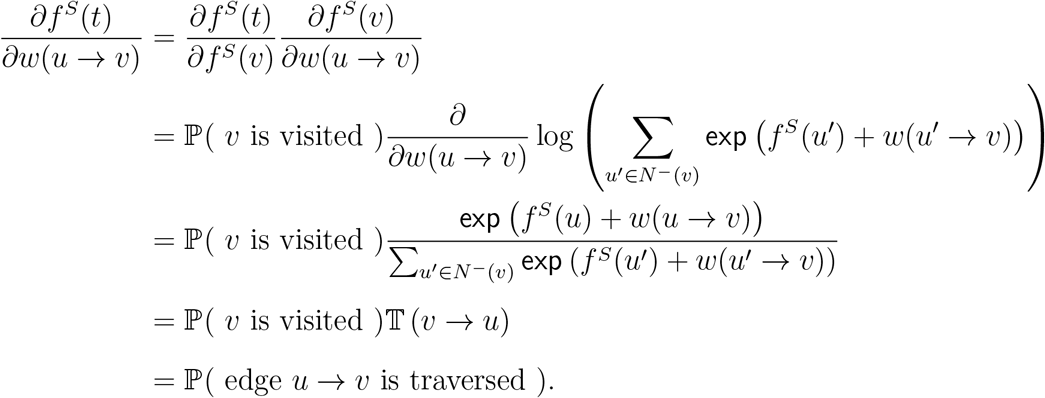

#### B. Difference in Needleman-Wunsch implementation of Morton et. al

The authors of [43] implement a differentiable version of the Needleman-Wunsch global alignment algorithm [46]. Their implementation differs from ours in how gaps are parameterized. Consequently, their output indicates where gaps or matches are likely, whereas our output expresses matches in an expected alignment.

The authors of [43] define

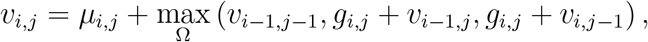

where *g*_*i,j*_ is the gap penalty for an insertion or deletion at *i* or *j, µ*_*i,j*_ is the alignment score for *X*_*i*_ and *Y*_*j*_, and max_Ω_(*x*) = log (_*i*_ exp (*x*_*i*_)) (see Appendix A of [43]). The values *v*_*i,j*_ are analogous to our definition *f* ^*S*^ on grid vertices (Equation (1)) with match scores *µ*_*i,j*_ = *a*_*i,j*_,

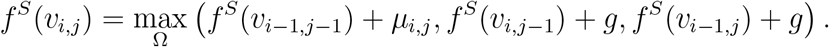

In the alignment graph for their formulation, gap edges have weight *µ*_*i,j*_ + *g*_*i,j*_. In our alignment graph, gap edges have weight *g*; the match score *µ*_*i,j*_ does not play a role, and our gap penalty is not position dependent.

Their code outputs the derivatives 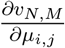. The derivative 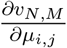 is high whenever the dominant alignment path uses an edge whose weight includes *µ*_*i,j*_; this includes the edges that corresponds to gaps. In contrast, in our formulation *a*_*i,j*_ = *µ*_*i,j*_ appears on the match edge only, and so 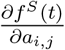 is high only when the dominant alignment path uses the edge corresponding to a match. Proposition 1 establishes that 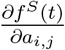 equal to the probability that *X*_*i*_ and *Y*_*j*_ are aligned, so our output is an expected alignment. Figure S2 establishes that this is not the case for the output of the Needleman-Wunsch implementation of [43].

**FIG. S2:**
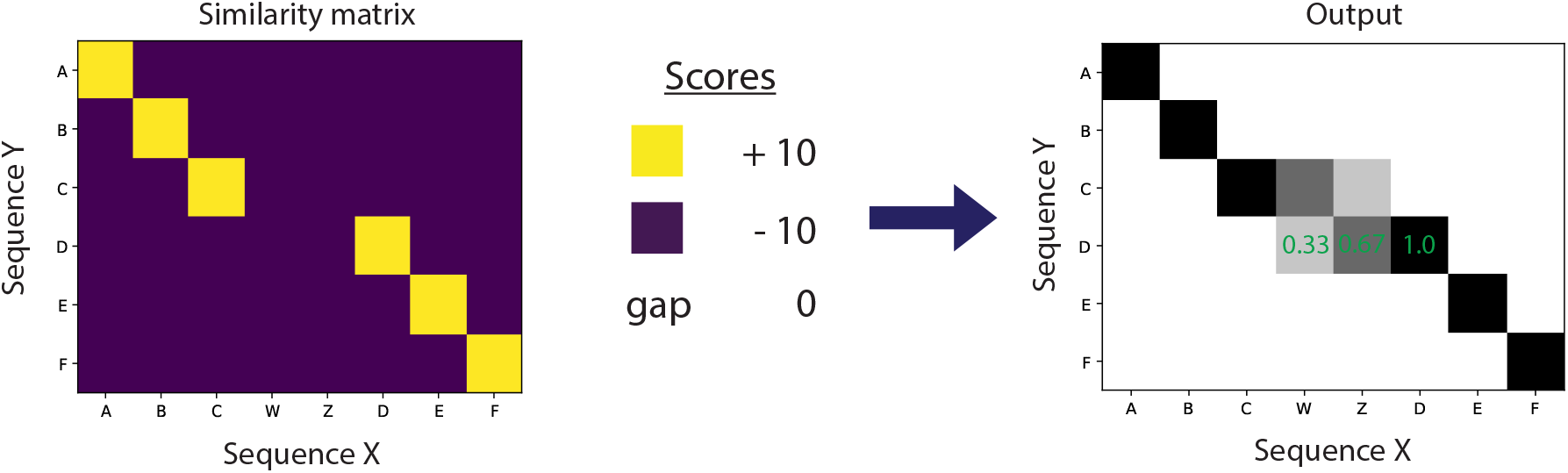
The output of the Needleman-Wunsch implementation of [43] is not an expected alignment. It is not the case that *Y*_4_ = *D* is aligned with *X*_4_ = *W* with probability 0.33, *X*_5_ = *Z* with probability 0.67, and *X*_6_ = *D* with probability 1.0 because in any alignment, *Y*_4_ can be aligned to at most one residue of sequence *X*.

#### C. Vectorization in our SSW implementation

Following the approach of Wozniak [64], we implement a version of smooth SmithWaterman where the values on the anti-diagonal are computed simultaneously. The vectorization speeds up our code substantially. In order to compute the final score *f* ^*S*^(*t*), we iteratively compute the scores of the grid vertices *f* ^*S*^(*v*_*i,j*_), which take as input the values *f* ^*S*^(*v*_*i−*1,*j*_), *f* ^*S*^(*v*_*i,j−*1_), and *f* ^*S*^(*v*_*i−*1,*j−*1_). In a simple implementation, a for loop over *i* and *j* is used to compute the values *f* ^*S*^(*v*_*i,j*_) (Figure S3a). To leverage vectorization, we instead compute the values *f* ^*S*^(*v*_*i,j*_) along each diagonal in tandem, i.e. all (*i, j*) such that *i* + *j* = *d*. To implement this, we rotate the matrix that stores the values *f* ^*S*^(*v*_*i,j*_) by 90 degrees so that each diagonal now corresponds to a row (see Figure S3b). In the rotated matrix, the values in a row *d* are a function of the values in rows *d −* 1 and *d −* 2, and therefore we can apply vectorization to quickly fill the matrix.

**FIG. S3:**
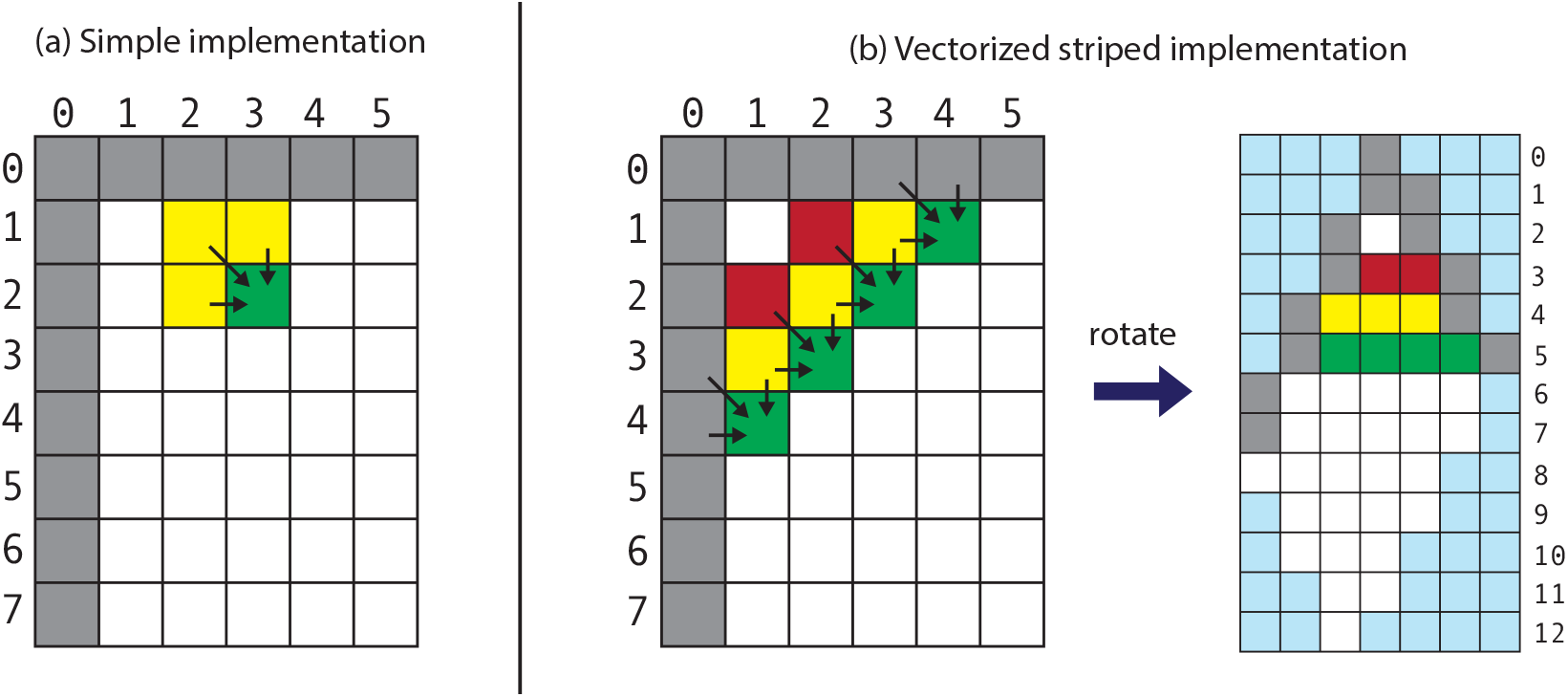
Vectorized implementation. (a) In a simple implementation, the value *f* ^*S*^(*v*_*i,j*_) are computed individually in a for loop over *i* and *j*. (b) In an anti-diagonal implementation, the values along each diagonal in the matrix are computed in tandem. We implement this with vectorization by rotating the matrix and computing the values in each row in tandem. The blue denotes meaningless positions in the rotated matrix that we set to *−∞*. This figure is inspired by Michael Brudno (University of Toronto).

#### D. SSW options

Our smooth Smith-Waterman implementation has the following four additional options.

##### a. Temperature parameter

The temperature parameter *T* controls the extent to which the probability distribution over alignments is concentrated on the most likely alignments; higher temperatures yield less concentrated alignments. We compute the smoothed score for the vertex *v* as

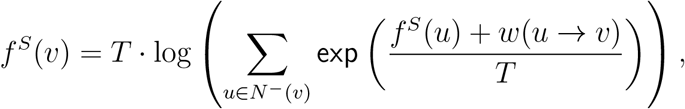

which matches Equation (1) at the default *T* = 1.

##### b. Affine gap penalty

The “affine gap” scoring scheme introduced to Smith-Waterman by [22] applies an “open” gap penalty to the first gap in a stretch of consecutive gaps and an “extend” gap penalty to each subsequent gap. The open gap penalty is usually larger than the extend penalty, thus penalizing length *L* gaps less severely than *L* separate single residue gaps.

To implement an affine gap penalty, we use a modified alignment graph with three sets \of grid vertices that keep track of whether the previous pair in the alignment was a gap or a match. Edges corresponding to the first gap in a stretch are weighted with the “open” gap penalty[48]. Figure S4a illustrates the incoming edges of the three grid vertices for (*i, j*). Paths corresponding to alignments with *x*_*i*_ and *y*_*j*_ matched pass through 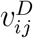, paths corresponding to alignments with a gap at *x*_*i*_ pass through 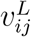, and paths corresponding to alignments with a gap at *y*_*j*_ pass through 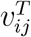. Storing three sets of grid vertices requires three times the memory used by the version with a linear gap penalty. For this reason we implemented SMURF with a linear gap penalty.

##### c. Restrict turns

Smooth Smith-Waterman is inherently biased towards alignments with an unmatched stretch of *X* followed directly by an unmatched stretch of *Y* over alignments with an equally long unmatched stretch in one sequence. Consider the example illustrated in Figure S4b where the highest scoring match states are depicted by bold black, light blue, and dark green lines. Suppose the match scores of the light blue and the dark green are identical. With a standard Smith-Waterman scoring scheme (no affine gap), the alignment containing the black and light blue segments has the same score as each alignment containing the black and dark green segments. However, there are more alignments that pass through the dark green segment. There are ten ways to align *ABC* and *V W* with no matches (the red, purple, orange, brown, and light green paths illustrate five such ways), but only one way to align *V WXY Z* with gaps (navy blue). Smooth Smith-Waterman will assign the same probability to each of these paths. However, since ten of the eleven paths go through the dark green segment, the expected alignment output by smooth Smith-Waterman will favor the dark green segment. This bias becomes more pronounced the longer the segments; there are 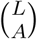 alignments of a sequence of length *L* and a sequence of length *L − A* with no matches.

To remove this bias, we implemented “restrict turns” option that forbids unmatched stretches in the *X* sequence from following an unmatched stretch in the *Y* sequence. To do so, we again use an alignment graph with three sets of grid vertices to keep track of the previous pair in the alignment. Removing the edge with the asterisk in Figure S4a, forbids transitions from an unmatched stretch in the *Y* sequence to an unmatched stretch in the *X* sequence. When implemented with this restrict turns option, smooth Smith-Waterman will find exactly one path through the dark green and black segments in Figure S4: the path highlighted in red. Due to the increased memory requirement of the restrict turn option, we did not utilize the option in SMURF.

##### d. Global Alignment

We also implement the Needleman-Wunsch algorithm, which outputs global alignments rather than local alignments.

**FIG. S4:**
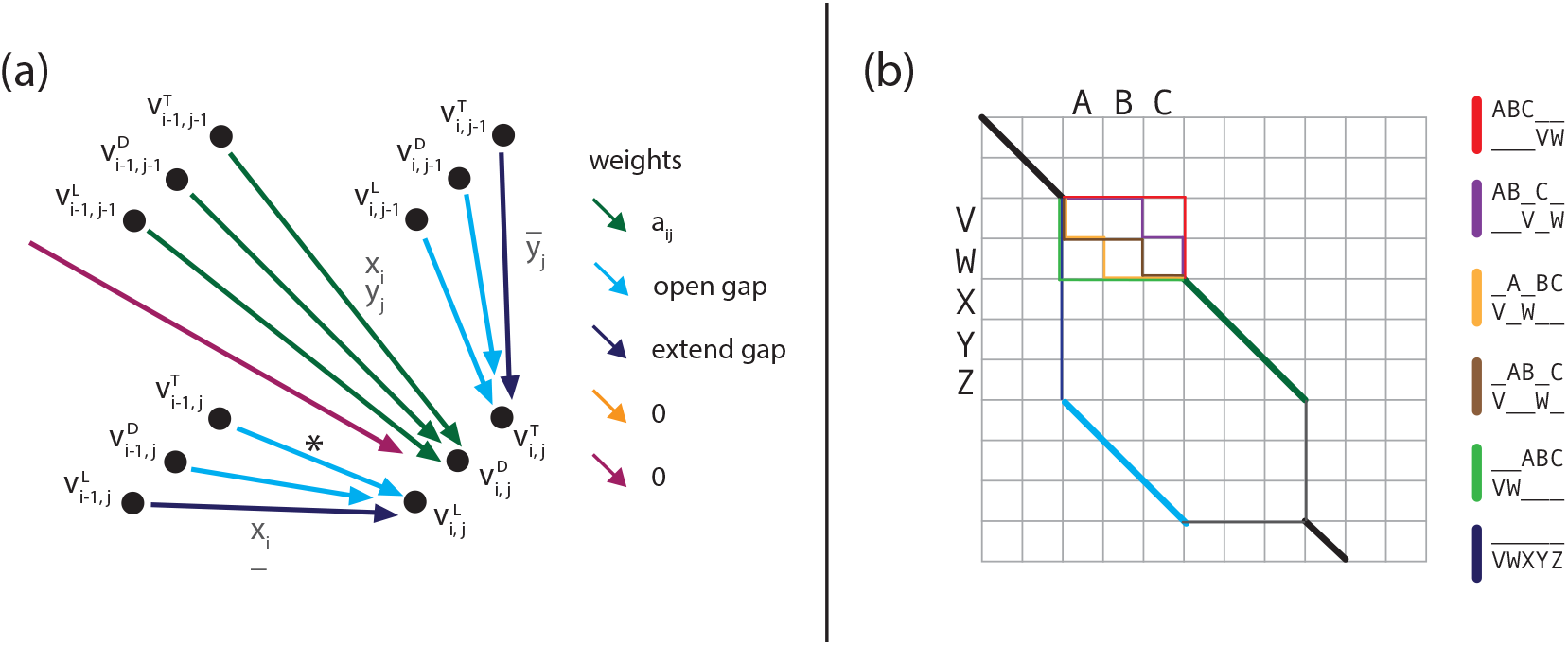
Algorithm modification for the affine gap penalty and restrict turns options. (a) The modification of the alignment graph from Figure 5 needed for the affine gap penalty. Incoming edges of the vertices 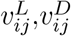, and 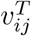 are illustrated. The colors of the edges indicate their weights. The grey labels describe the corresponding aligned pair for each group of edges. The red edge is incoming from the source vertex *s*. There is an outgoing edge from 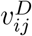 to the sink *t* for all *i, j ≥* 1 (not pictured). The edge marked with an asterisk is removed under the “restrict turns” option. (b) Without the restrict turns option, there ten paths containing both black segments and dark green segment. The red, purple, orange, brown, and light green illustrate five of these paths. There is only one path that contains both black segments and light blue segment, as depicted in navy blue. The sub-alignments corresponding to the colored segments are written on the right. With the restrict turns option the purple, orange, brown, and green paths are not valid.

**FIG. S5:**
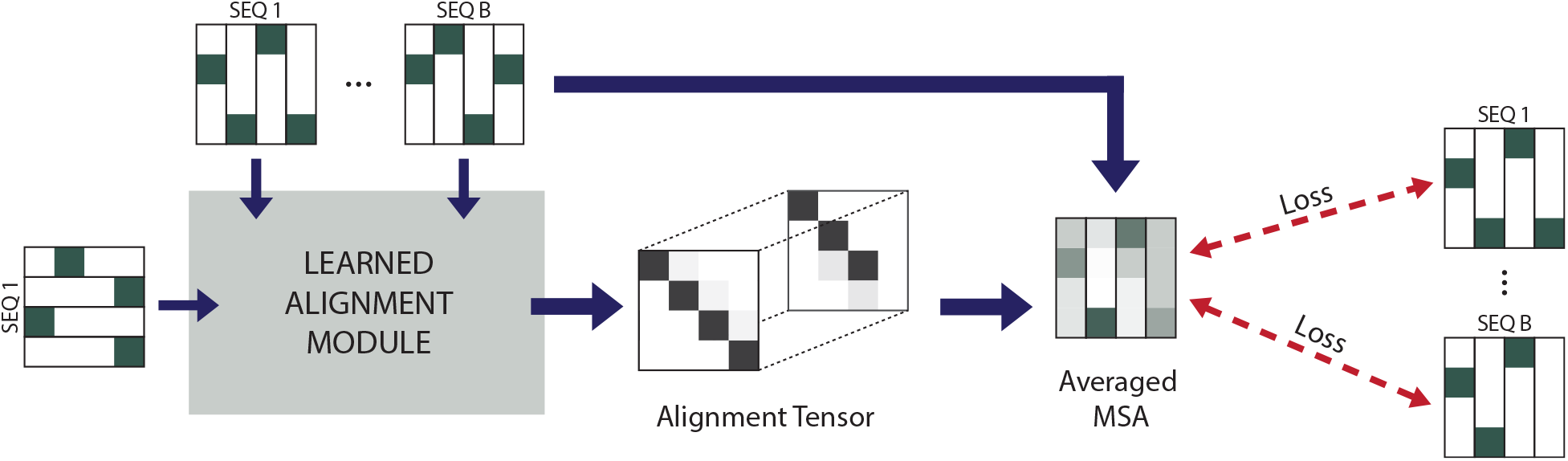
BasicAlign. An alignment is computed with the learned alignment module (Figure 1), and the corresponding MSA is averaged. Squared loss (Equation (4)) is computed between the averaged MSA and the one-hot encoding of the aligned input sequences.

**FIG. S6:**
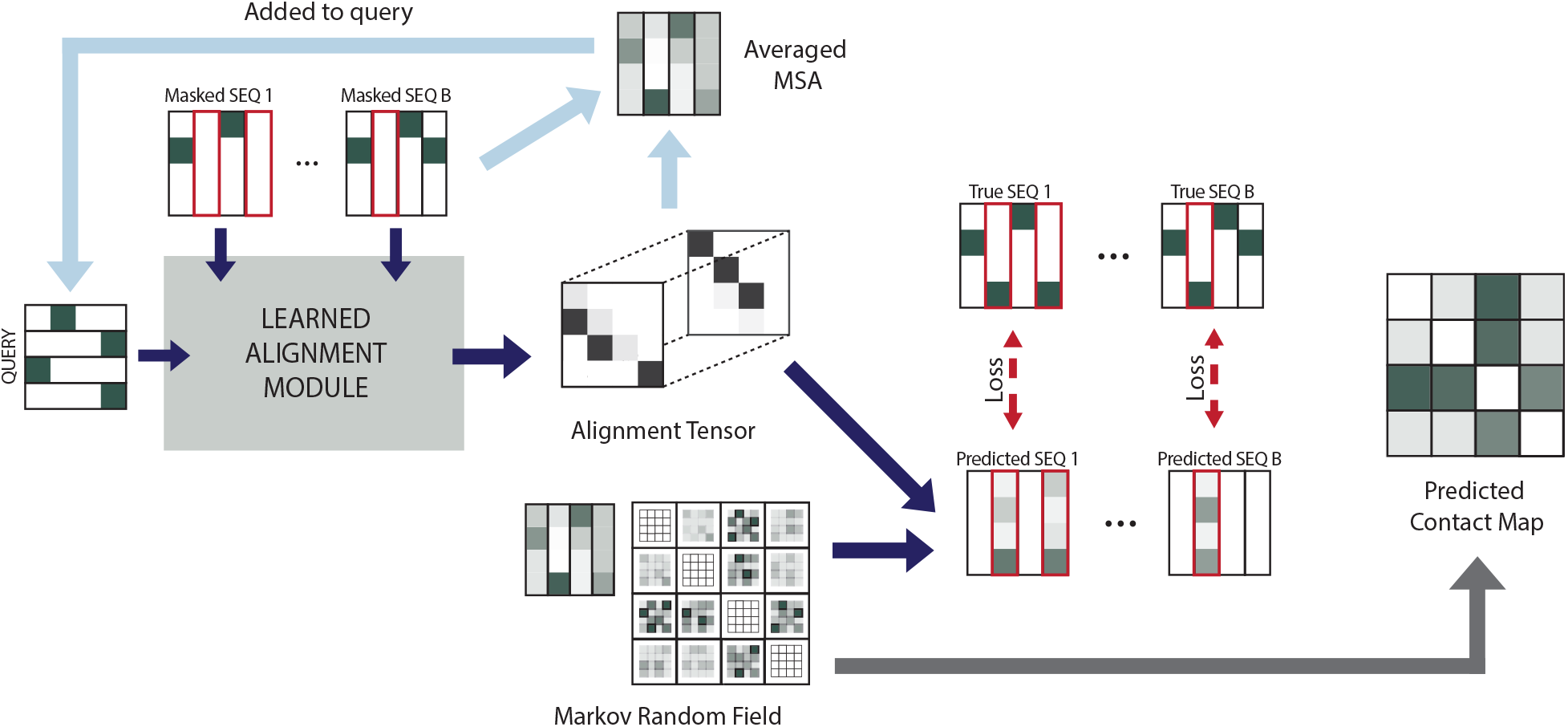
TrainMRF. Random positions in the input sequences are masked, then aligned with the LAM (Figure 1). A prediction for the masked positions is computed from the MRF parameters according to Equation (5). The network is trained with cross entropy loss given by Equation (6). The light blue arrows illustrate the update to the query that occurs between iterations of training; the query is a weighted average of the one-hot query sequence and a running average of the MSAs computed in previous iterations, see Equation (7). The grey arrow depicts the extraction of the contact map from the MRF matrix *w* at the end of training, as described in Equation (8).

**FIG. S7:**
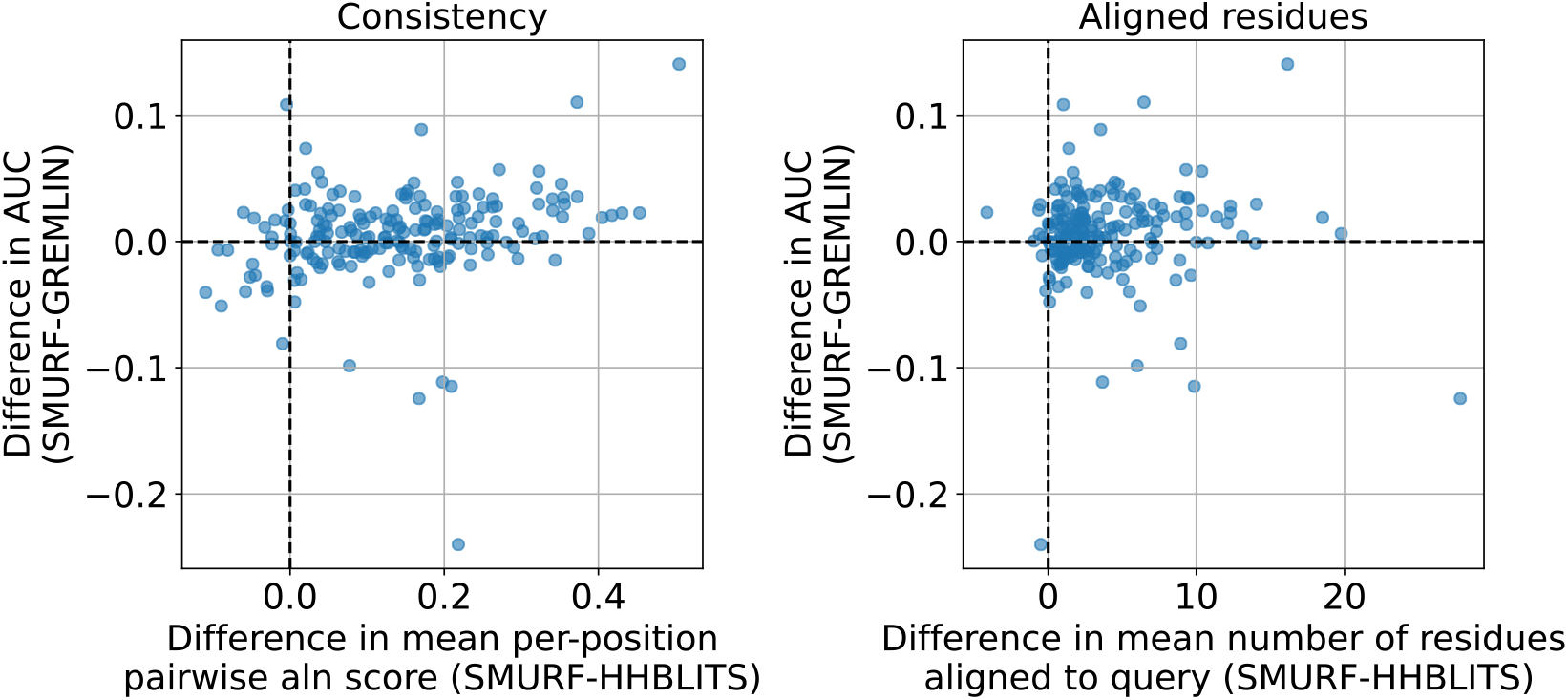
SMURF-learned alignments are more consistent and have more residues aligned to the query in comparison to HHBlits alignments. Left: The BLOSUM pairwise alignment scores are on average higher for SMURF MSAs as compared to HHBlits MSAs. There is a postive correlation between an increase in pairwise alignment score and the improvement of SMURF over GREMLIN contact accuracy prediction. BLOSUM scores were computed only over positions that correspond to a residue in query sequence and used an affine gap penalty with open penalty *−*11 and extend penalty *−*1. Right: SMURF MSAs tend to have more positions aligned to the query as compared to HHBlits MSAs. This quantity does not appear correlated with the relative performance of SMURF over GREMLIN. Both plots were generated from a random sample of 50 sequences from each alignment (out of 1024 sequences).

**FIG. S8:**
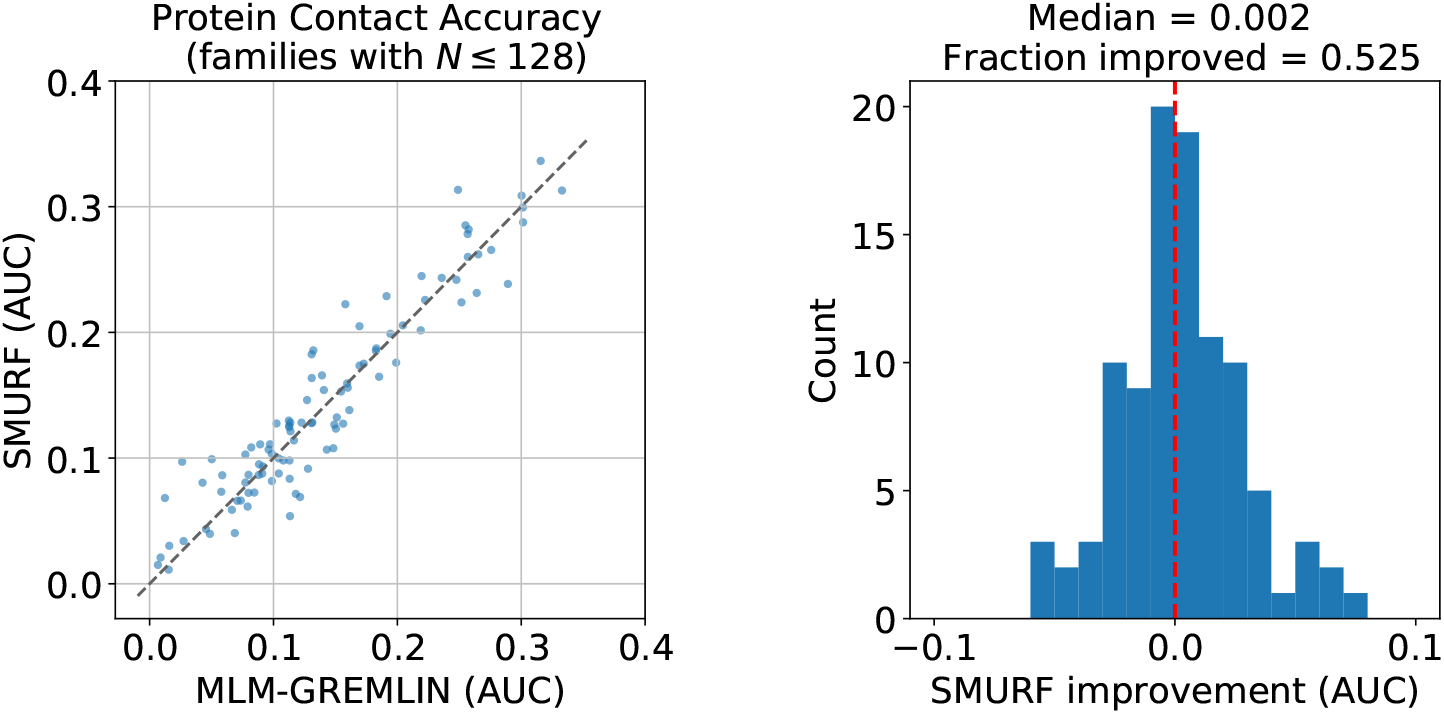
SMURF performance on 99 protein families from [4] with at most 128 sequences. Left: Scatter plot of the AUC of the top L predicted contacts for SMURF versus MLM-GREMLIN. Right: Histogram of the difference in AUC between SMURF and MLM-GREMLIN.

### S2. SUPPLEMENTARY NOTE: FURTHER ANALYSIS OF EXAMPLE SMURF PREDICTIONS

#### A. RNA contact prediction

By comparing the positive predictive value (PPV) for different numbers of predicted contacts, we see that SMURF consistently yields a higher PPV for RFAM family RF00167 (Figure 2b). For RF00010, it starts off higher but then drops off faster, leading to a lower overall AUC. Upon a visual inspection of the contact predictions, MLM-GREMLIN evidently generates more false positive predictions in seemingly random locations. On the other hand, SMURF largely resolves this issue, even for RF00010, presumably as a result of a better alignment. Interestingly, SMURF’s lower AUC for RF00010 can be attributed to a concentration of false positive predictions near the 5’ and 3’ ends. It remains unclear whether these represent a coevolution-based structural element that was not present in the specific RNA sequence deposited in PDB or whether these arise from artifacts of the learned alignment.

#### B. Protein contact prediction and alignments

Next, we investigated the contact predictions and alignments produced by SMURF. Figure S9 and Figure S10 illustrate the contact predictions, corresponding positive predictive value (PPV) plots, and alignments for the three families that improved the most and least (respectively) under SMURF as compared to MLM-GREMLIN. The poor performance of SMURF on 3LF9A can be attributed to the misalignment of the first *≈* 25 residues of many sequences (including the one illustrated) to positions *≈* 75 to 100 of the reference rather than to the first 25 positions of the reference. This is likely because the gap penalty for leaving positions *≈* 25 to 75 unaligned outweighs the benefit of aligning to beginning of the reference. Since our code computes a local alignment, there is no penalty for leaving positions at the beginning of the reference unaligned. Perhaps using our implementation of Smith-Waterman with an affine gap penalty would lead the network to learn a less severe penalty for long gaps and arrive at correct alignment. For the most improved families, we see that SMURF tends to predict fewer false positive predictions in seemingly random positions, as observed for RNA.

**FIG. S9:**
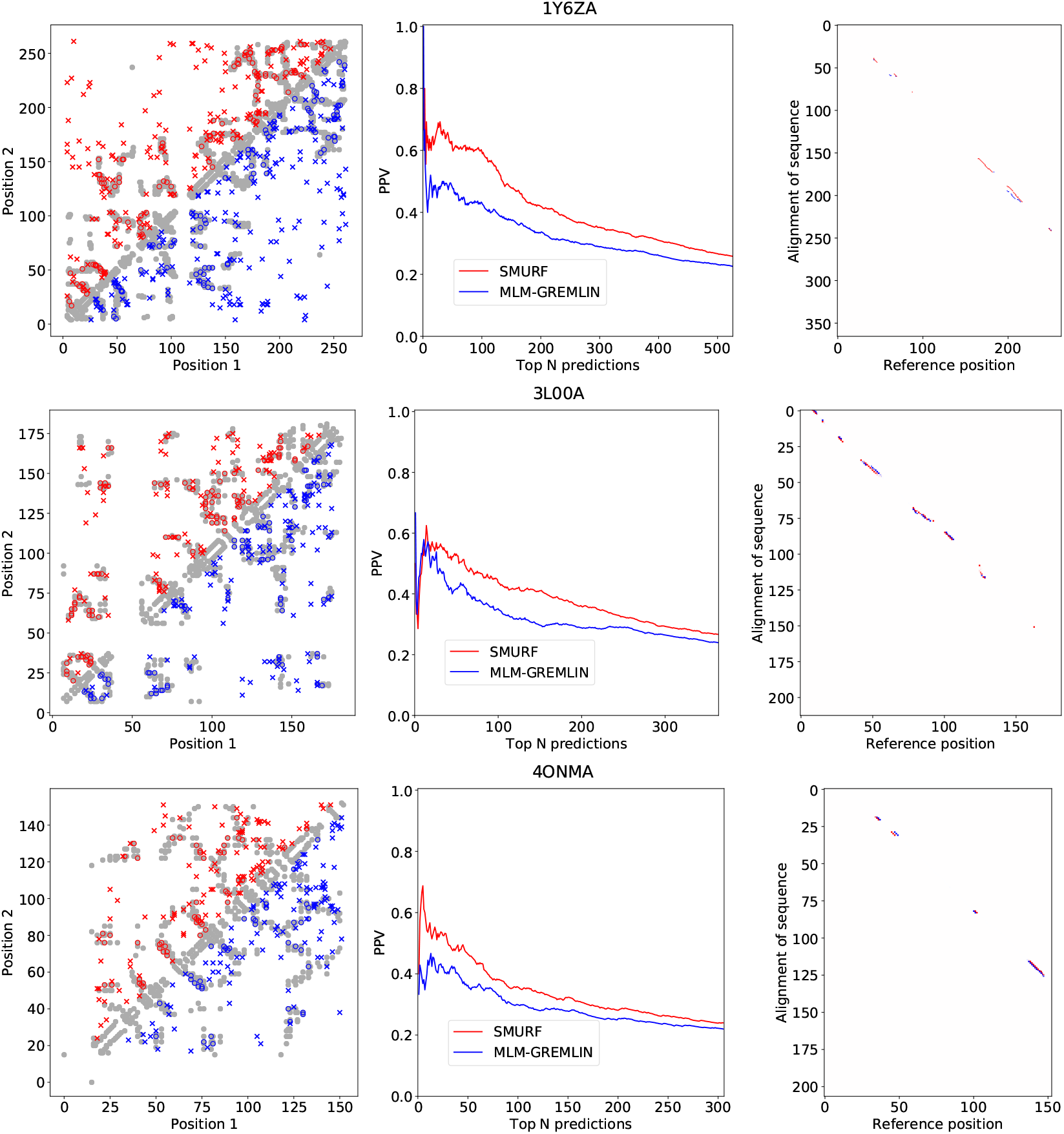
Contact predictions and alignments for the three most improved protein families. Left: Comparison of contact predictions between SMURF (red) and MLM-GREMLIN (blue). Gray dots represent PDB-derived contacts, circles represent a true positive prediction, and x represents a false positive prediction. Middle: The positive predictive value (PPV) for different numbers of top *N* predicted contacts, with *N* ranging from 0 to 2*L*. Right: Comparison of the alignment of a random sequence in the family to the reference sequence. Red indicates aligned pairs that appear in the SMURF alignment, but do not appear in the given alignment. Blue indicate aligned pairs that appear in the given alignment, but do not appear the alignment found by SMURF.

**FIG. S10:**
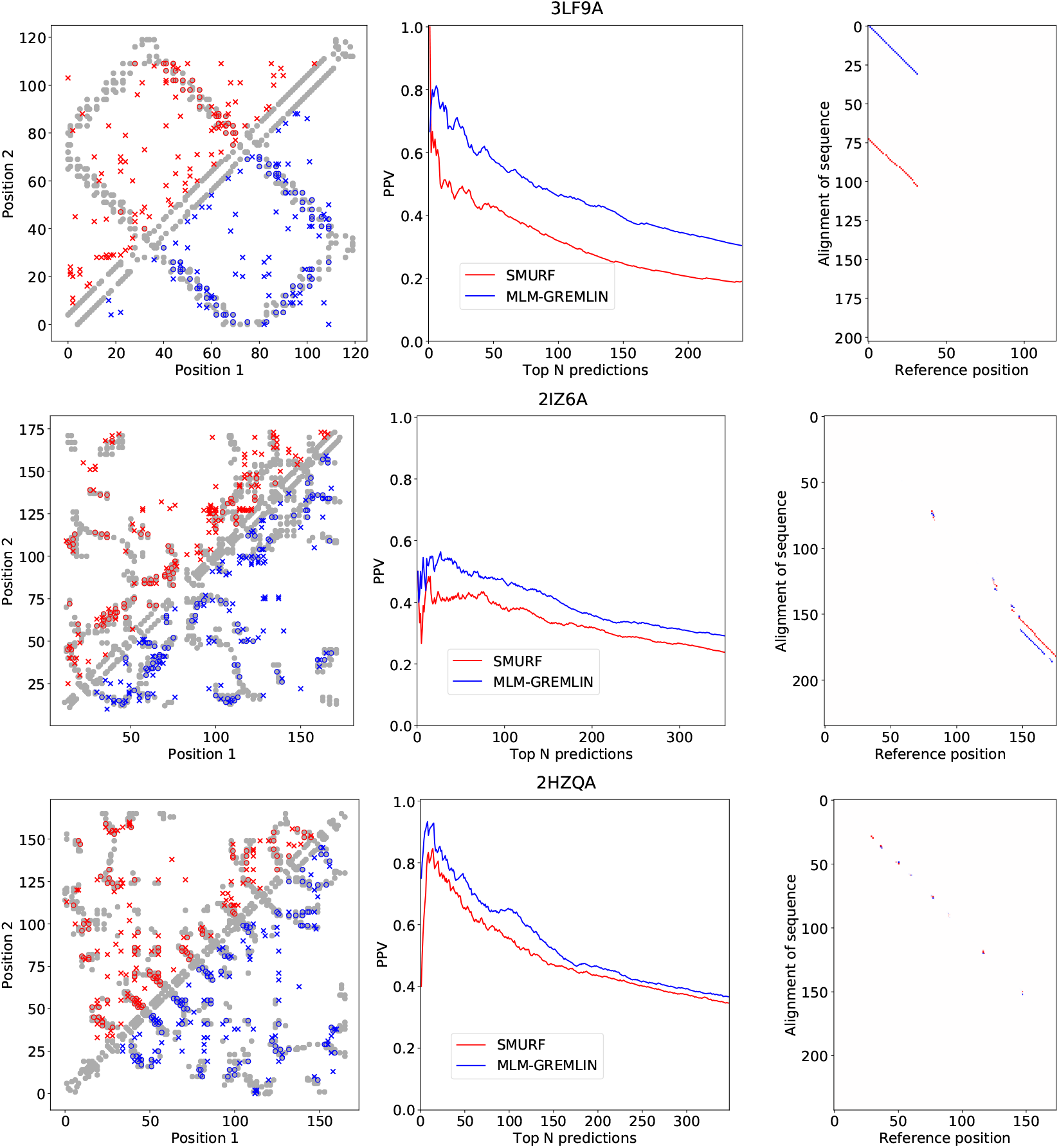
Contact predictions and alignments for the three worst performing protein families (as compared to MLM-GREMLIN). Left: Comparison of contact predictions between SMURF (red) and MLM-GREMLIN (blue). Gray dots represent PDB-derived contacts, circles represent a true positive prediction, and x represents a false positive prediction. Middle: The positive predictive value (PPV) for different numbers of top *N* predicted contacts, with *N* ranging from 0 to 2*L*. Right: Comparison of the alignment of a random sequence in the family to the reference sequence. Red indicates aligned pairs that appear in the SMURF alignment, but do not appear in the given alignment. Blue indicate aligned pairs that appear in the given alignment, but do not appear the alignment found by SMURF.

**FIG. S11:**
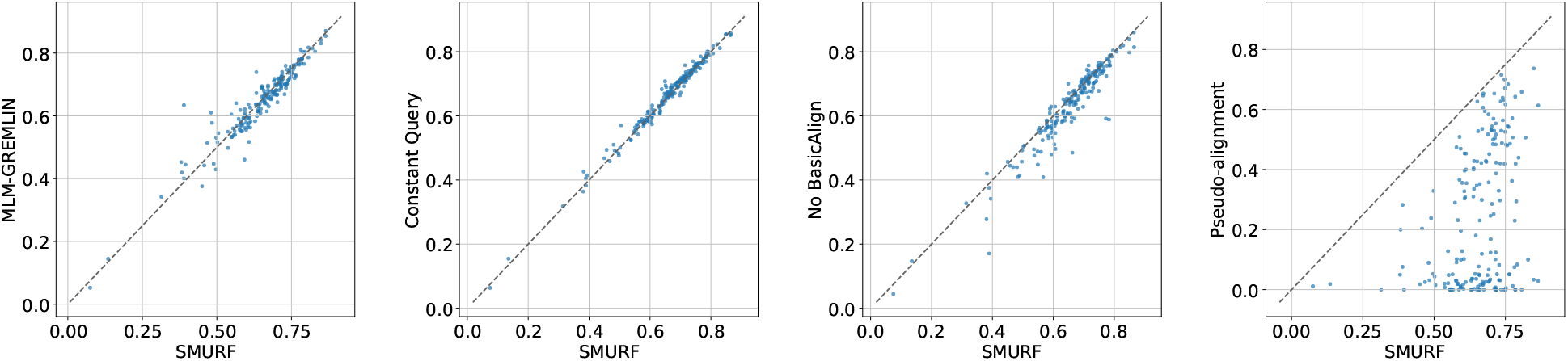
Ablation results. Contact AUC for SMURF versus ablated methods. Each point represents one family in the test set. In “Constant Query,” we did not update the the query with the averaged MSA between iterations (as depicted by light blue arrows in Fig. S6). In “No BasicAlign,” the convolutions were not initialized with BasicAlign, and instead TrainMRF was run for 4000 iterations. In “pseudo-alignment,” we replaced Smith-Waterman with a pseudo-alignment obtained by taking the softmax of the similarity matrix row-wise and column-wise, multiplying the resultant matrices, and taking the square root (similar to [7]).

**FIG. S12:**
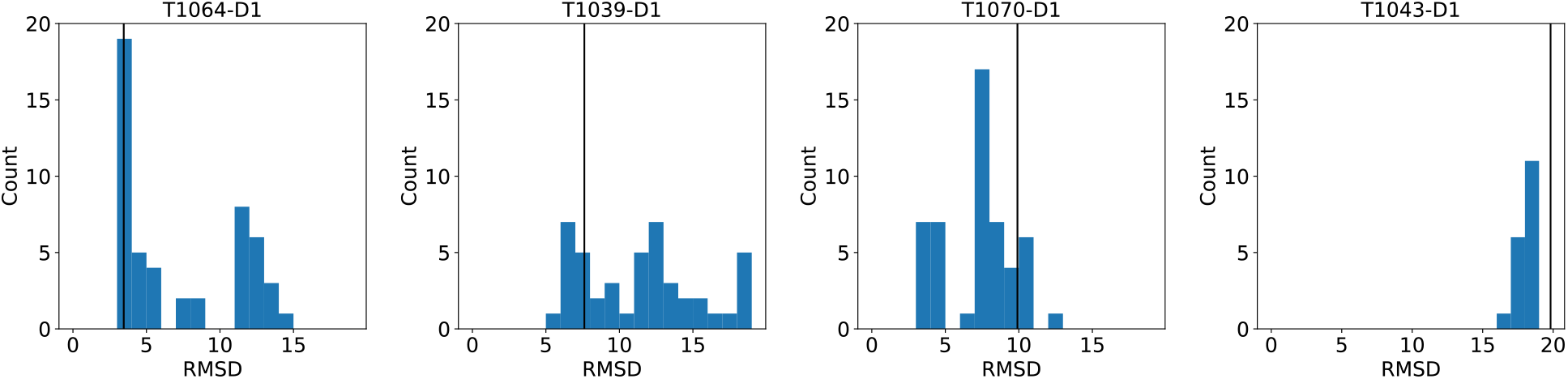
Sensitivity of AlphaFold predictions to random masking. By default, a random mask is used when AlphaFold makes a structure prediction [30]. The distribution of RMSD of AlphaFold predictions for MMSeqs2 MSAs with different random seeds used for the masks. The black line shows the RMSD of the prediction without the mask.

**FIG. S13:**
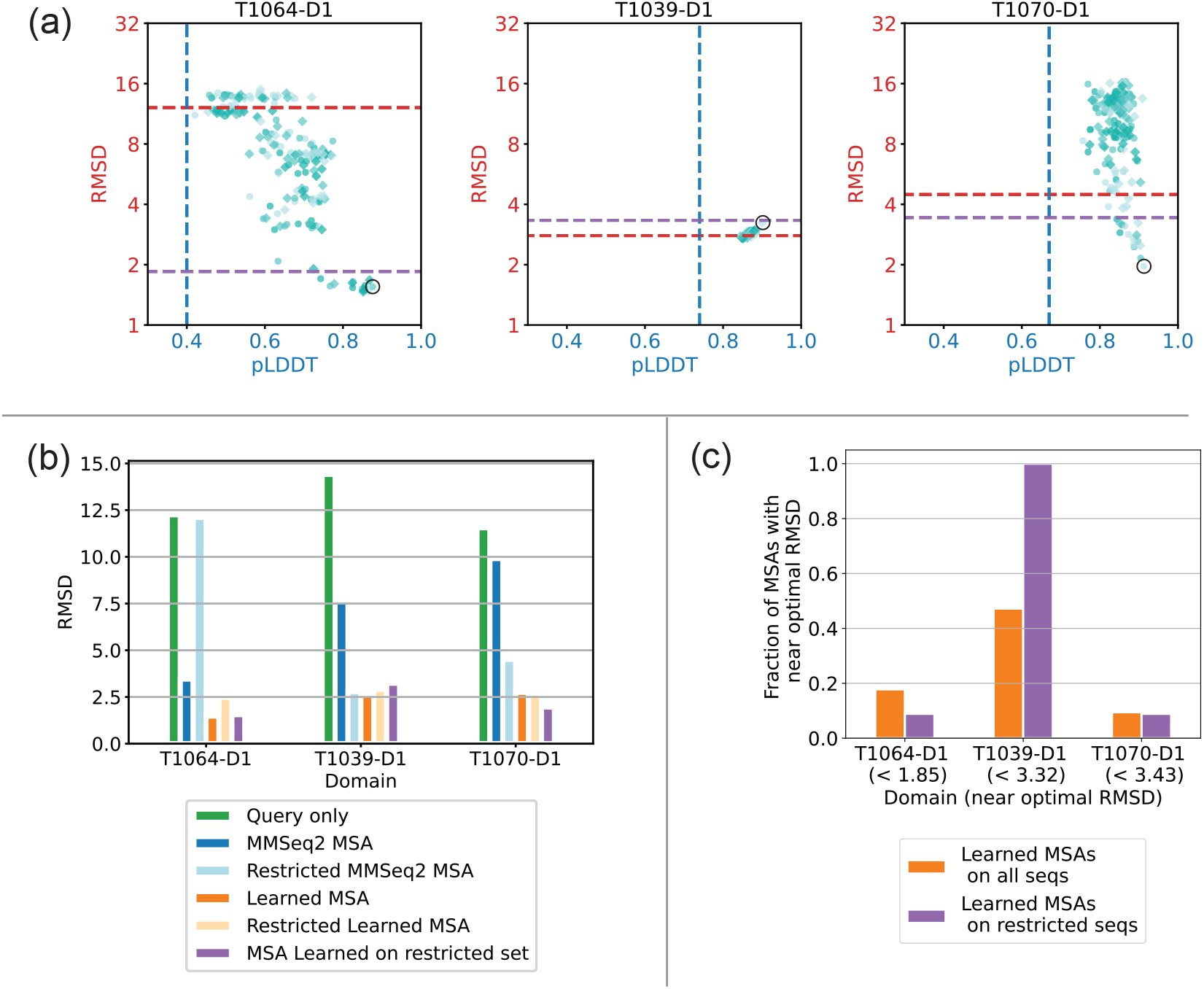
AlphaFold + LAM optimization on families with distant sequences removed. (a) Analogous plot to Fig. 3 for LAM + AF experiment with the distant sequences removed. The dotted blue and red lines show the pLDDT and RMSD of the prediction using the MSA from MMseqs2 with the distant sequences removed. The purple line indicates the definition of “near-optimal” and is 1.25 times the RMSD of the prediction for the “Learned MSA” found in Fig. 3 or 4. We selected the circled point maximizing the confidence (pLDDT) as our “MSA Learned on restricted set.” (b) A comparison of the RMSD for various tested MSAs, by domain. (c) Fraction of MSAs learned that yielded predictions with “near optimal” structure.

